# Reinforcement Learning Reveals Spatial Strategies for Slowing Biological Invasions

**DOI:** 10.64898/2026.10.28.762500

**Authors:** Jean-Nicolas Lang, Adam Lampert

## Abstract

Managing biological invasions requires deciding where, when, and how to act while populations continue to grow and spread. Invasive species threaten ecosystems and livelihoods, but limited management resources mean that managers cannot treat every affected area. This study asks how control efforts can be directed across a landscape to slow an invasion while keeping costs low. We develop a reinforcement-learning framework that can use broader spatial context to control biological invasions, using the spongy moth as a case study. The stochastic simulator captures host-dependent growth, short- and long-distance dispersal, density-dependent mating, and the effects of multiple treatments. The reinforcement-learning strategies are compared with benchmarks that are based on local treatment rules. We find that reinforcement learning achieves the lowest mean cost, with its advantage widening as management complexity increases, such as with simultaneous advancing-front control, nascent-colony management, and established-colony containment on heterogeneous-host landscapes. The reinforcement-learning strategies exhibit highly context-dependent treatment behavior, adjusting treatment choices as invasion dynamics evolve and coordinating interventions across long distances, all of which is learned from scratch.

## 1 Introduction

Artificial intelligence has made substantial advances in ecological monitoring and prediction. Machine-learning methods are increasingly used to identify species, map risk, forecast ecological change, and extract patterns from large ecological data sets^1;2^. Effective ecological management requires a further step, namely deciding where, when, and how to intervene while the system continues to change. This gap is evident in the management of invasive species, which threaten biodiversity, ecosystem functioning, agriculture, forestry, and human well-being^3–5^, while global economic costs of biological invasions have increased over time^6^. Because managers cannot treat every invaded or threatened location indefinitely, effective control requires identifying interventions that limit future spread while keeping management costs low.

Predicting where an invasive population will be abundant does not, by itself, reveal where treatment will be most valuable. For example, a small satellite colony beyond the main invaded area may become a new source of spread if left untreated, whereas a dense population behind the main invasion front may contribute relatively little to further expansion (Figure 1 **A**)^7–9^. This distinction between current abundance and future spread has long informed invasion ecology and bioeconomic theory. Bioeconomic models show that choices about prevention, detection, and control depend on the risk of new introductions, spread dynamics, management costs, and damages^10–13^. Studies that account for spatial structure further show that treatment priorities depend on invasion-front position and the potential for future spread^14–17^. Spatial models also identify priority sites in dispersal networks^18^ and guide the design of buffer zones around conservation assets^19^. However, translating these insights into repeated treatment decisions across a large landscape remains computationally demanding.

**Figure 1.**
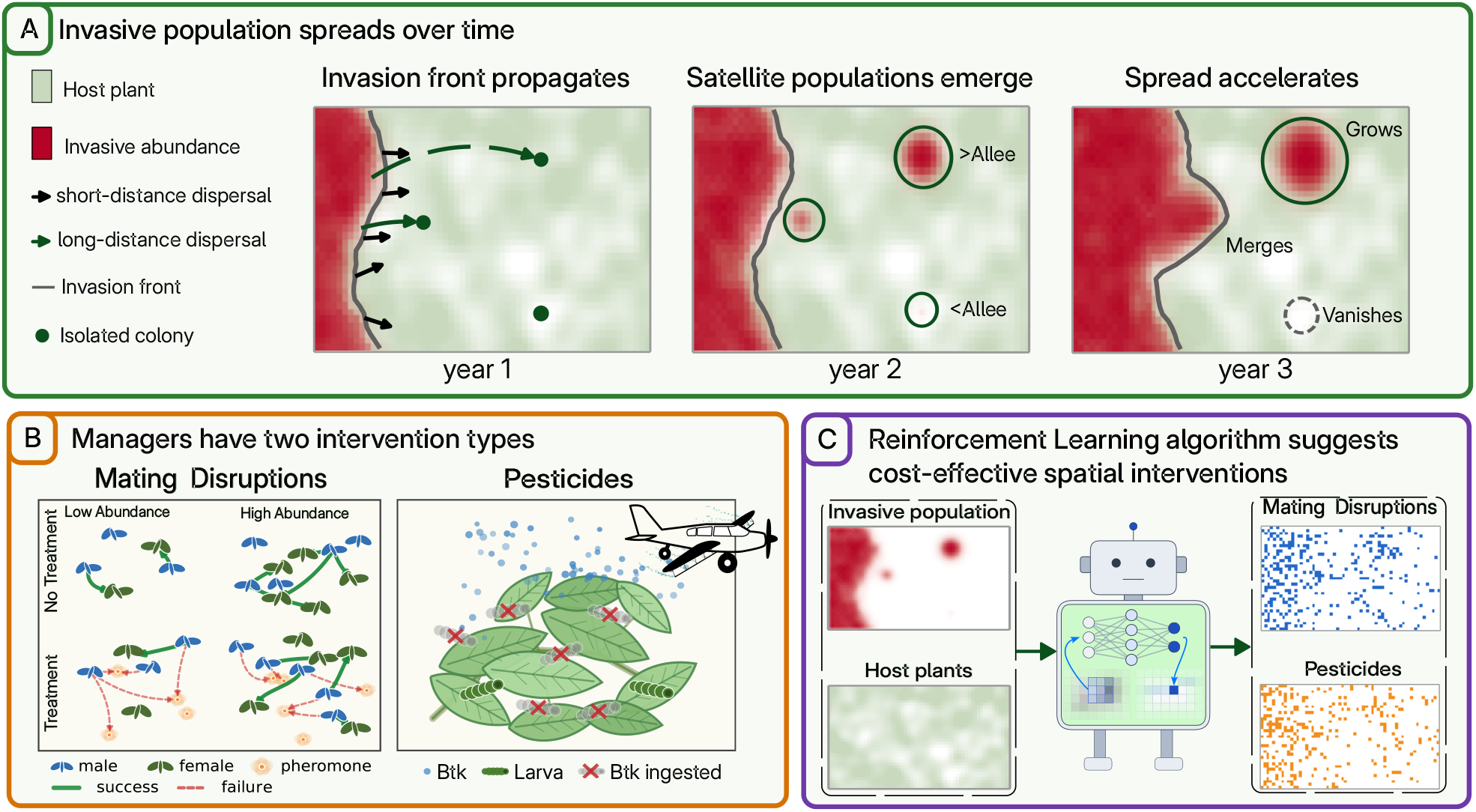
Overview of invasion dynamics, treatment mechanisms and optimization approach. **(A)** The population evolves through growth and dispersal processes. Spread combines local movement with occasional long-distance dispersal that can establish satellite colonies. Small satellites may collapse due to Allee effects, whereas established satellites can accelerate range expansion. **(B)** The manager seeks to slow the spread cost-effectively by applying mating disruption and pesticides. Mating disruption reduces mate-finding success at low population densities, while pesticide application directly suppresses local abundance at higher cost. **(C)** We train an agent with a novel reinforcement-learning algorithm to select cost-effective spatial interventions based on the observed state. Specifically, the goal is to decide, at each time step, where to apply treatment across the landscape. The algorithm is specifically designed for large landscapes, where discrete treatment choices across cells create a combinatorial action space (see Methods).

Bioeconomic models formulate such management problems in terms of the long-term costs and benefits of interventions^16;20^. Dynamic optimization provides one way to solve these models by evaluating decisions and their consequences over time ^21^. For example, simulator-based planning has been used to manage invasive tamarisk in river networks^22^, while graph-based Markov decision processes have enabled near-optimal simultaneous invasive-species control across ecological networks^23^. However, finding optimal solutions becomes more complicated when the problem is spatially explicit and stochastic^24^. Decisions must then be coordinated across many locations while accounting for uncertain spread, variation in habitat, multiple treatment options, and limited resources. Present interventions also change future spread and, therefore, the decisions that will be needed later. Depending on the method, obtaining tractable solutions may require reducing the spatial domain, simplifying the action set, or assuming a restricted class of policies^25^. These simplifications are useful, but they can limit the spatial strategies explored or the ecological detail retained. The challenge is to identify effective strategies while retaining the features relevant to future spread.

Reinforcement learning (RL) offers an approximate approach to solving these bioeconomic management problems. An RL agent learns through interactions with a dynamic system, using the long-term consequences of its actions to improve its decisions ^26;27^. The agent learns a policy that specifies which management actions to take based on the available information about the system’s state. RL has been applied to natural-resource management^28^, invasive-species management^29^, and crop pest and disease management^30^. Like adaptive management, RL emphasizes learning through experience^31^. Specifically, deep RL can process high-dimensional observations^32^ and has been applied to conservation planning^33;34^ and connectivity planning^35^. At the same time, applying RL to spatial invasion control, where treatment decisions must be coordinated across a landscape while accounting for uncertain spread, presents several methodological challenges. Choosing a treatment type at each location creates a combinatorial action space: the number of possible landscape-wide treatment combinations grows exponentially with the number of locations^36;37^. Evaluating every combination becomes impractical^38^, yet basing treatment only on local conditions can overlook how interventions jointly affect future spread. Building on these approaches, the challenge addressed here is to learn treatment maps without exhaustive search and assess whether policies trained on smaller landscapes remain effective on larger, heterogeneous landscapes with stochastic spread and overlapping management tasks.

Here, we develop a deep RL framework for cost-effective invasion management across large landscapes, using the spongy moth (*Lymantria dispar*) in North America as a case study (Figure 1 **B,C**). Spongy moth is a major invasive forest pest^39^ and the target of the long-running Slow the Spread program. The program uses surveillance and treatment across a broad invasion front to delay range expansion^40;41^. The system illustrates a broader management challenge: deciding where treatment will do the most to slow future spread, not just where populations are most abundant. Managers must balance slowing an advancing invasion front with stopping new colonies farther away, while responding to differences in host trees, colony survival, and treatment effects. Colony survival, for example, depends on both moth density and the area a colony occupies^42^. Recent models show that suppression zones, host gaps, and spatial treatment allocation can change management cost-effectiveness^43–45^. Building on these insights, we examine whether RL can learn to coordinate treatments using spatial context on a large, two-dimensional landscape, where dynamics are stochastic and interactions among sub-colonies can shape future spread. Our framework uses a learned evaluator to improve proposed treatment maps and trains a policy to reproduce the improved choices, without evaluating every possible combination. In our RL framework, strategies learned on smaller landscapes remain effective on larger ones without further training and achieve lower mean combined long-term costs of treatment and invasion damage than optimized local treatment rules. The learned policies adjust treatment priorities to host availability and each colony’s position relative to the invasion front. The central contribution is a demonstration that RL can learn and combine spatial treatment strategies in large-scale, ecologically complex simulations without those strategies being prescribed in advance.

## 2 Results

### Model and study design

We study spatial invasion control with a stochastic model of spongy moth spread on a two-dimensional landscape (Figure 1 **A**). Each annual time step represents one life cycle. The state in each cell is represented by continuous measures of adult abundance, host-plant abundance, and invasion persistence (see Methods). Local population growth depends on host-plant density, mortality, and density-dependent mating and reproduction. This growth is characterized by an Allee effect in which small, low-density colonies may fail to persist, whereas larger colonies are more likely to grow ^46^ (Figure S2). In each annual time step, local growth is followed by stochastic dispersal between cells, including both short-distance spread and rarer long-distance spread, which can establish satellite colonies ahead of the main front^47^. The complete dynamic model is described in detail in Methods. Our simulations yield spread rates under intensive mating disruption and without intervention that are comparable to those reported in studies of the Slow the Spread program^48;49^, when each cell represents approximately 12 km × 12 km (Figure S1).

Managing a spatial invasion requires repeated decisions about where and how to intervene as the population grows and disperses across the landscape (Figure 1 **B**). We consider a single manager with two available intervention methods: mating disruption and pesticide application. Mating disruption reduces mating success and is most effective at low population density. Pesticide is more expensive but reduces population size at both low and high densities. Each year, the manager observes the full state and decides for each and every cell whether to apply mating disruption, pesticide, both, or neither. The manager’s objective is to minimize the expected discounted cumulative cost, incorporating both treatment expenditures and the damage caused by the invasion.

The complex dynamic model combined with the large state and action spaces necessitates the use of approximation techniques. Here we use deep reinforcement learning (RL) (Figure 1 **C**) to optimize the treatment strategy. Our aim is to train a generalist agent capable of managing previously unseen invasion scenarios across large spatial domains.

Large but finite discrete action spaces are already very challenging and call for extensions to standard RL methods^38;50^. In real-world management, the spatial extent of the management problem may vary over time. This motivates us to make both the agent and its training process agnostic to landscape size, so that the same agent can, in principle, be trained on and applied to problems of arbitrary size. The latter is enabled by a fully convolutional parameterization of the final policy. This size-agnostic training approach must accommodate a discrete action space whose size is not fixed a priori. To this end, we develop a variant of an actor–critic algorithm^51^ in which all components are agnostic to action-space size. Specifically, the actor proposes an initial action which the critic then refines^38^. In our case this refinement is performed through a gradient-based search (Figure 6 **B**). The actor simultaneously serves as an apprentice^52;53^, learning through imitation to reproduce these improved actions. To enable generalization across invasion conditions, our agent is exposed to smaller-scale training tasks that are procedurally generated and vary in terms of the initial invasion and host-plant landscapes. The training set comprises five tasks on a 10 ×40 grid: three broad-front invasions under uniform host cover, uniform host cover with a gap, or heterogeneous host cover, and two colony-management tasks involving one or two isolated colonies under uniform host cover. These tasks can be thought of as snapshots of the target large-scale landscape, taken at different times and in different locations, thereby exposing the agent to the varying shapes and progress of the invasion. Details of the training tasks and the RL algorithm are provided in Methods.

We evaluate the agent using independent simulations of the training tasks. The larger-grid setups referred to as *generalization tasks* retain the dynamics used during training, whereas the more challenging *late-detection generalization tasks* extend them with rare stochastic events that seed sizable off-front populations. These events mimic the sudden discovery of established colonies; once introduced, the colonies are fully observed by the manager. The resulting management problem therefore requires the agent to combine front control that slows the advancing front with off-front management that prevents new satellite colonies from establishing and contains colonies that are already established. The generalist RL policy is evaluated on these larger-grid tasks without further training. For comparison, we also retrain the generalist agent separately on each larger-grid task, producing task-specific RL agents (see Methods). To assess sensitivity to dispersal assumptions and invasion-damage costs, we compare performance under two long-distance dispersal kernels and two damage-cost levels, with agents trained separately for each combination.

### RL generalizes to larger grids with overlapping management tasks

Our central result is that the generalist RL policy adapts its treatment decisions to spatial context beyond local abundance and gradient. It generalizes to larger domains with overlapping management problems, and outperforms optimized local decision rules. The advantage of RL is largest in complex management settings, where treatment must address several spatial management problems simultaneously. For example, the RL agent treats small off-front satellite colonies more aggressively than segments of the main front with similar abundance and gradient magnitude (Figure 2). In particular, once an off-front satellite population is established and expanding, the RL agent suppresses it by applying pesticides within the satellite population and mating disruption around it. Management of the main invasion front is likewise context dependent. Stochastic spread and host heterogeneity naturally cause the front to deviate from a straight line. The generalist RL policy counteracts these distortions by concentrating treatment on sections furthest ahead while reducing treatment where the front lags behind or crosses areas with low host availability. The policy also reallocates treatment as sizable satellite colonies merge with the front, allowing the resulting bulges in the invasion front to contract gradually. These responses indicate that the policy adjusts treatment in one part of the landscape in response to invasion dynamics elsewhere.

**Figure 2.**
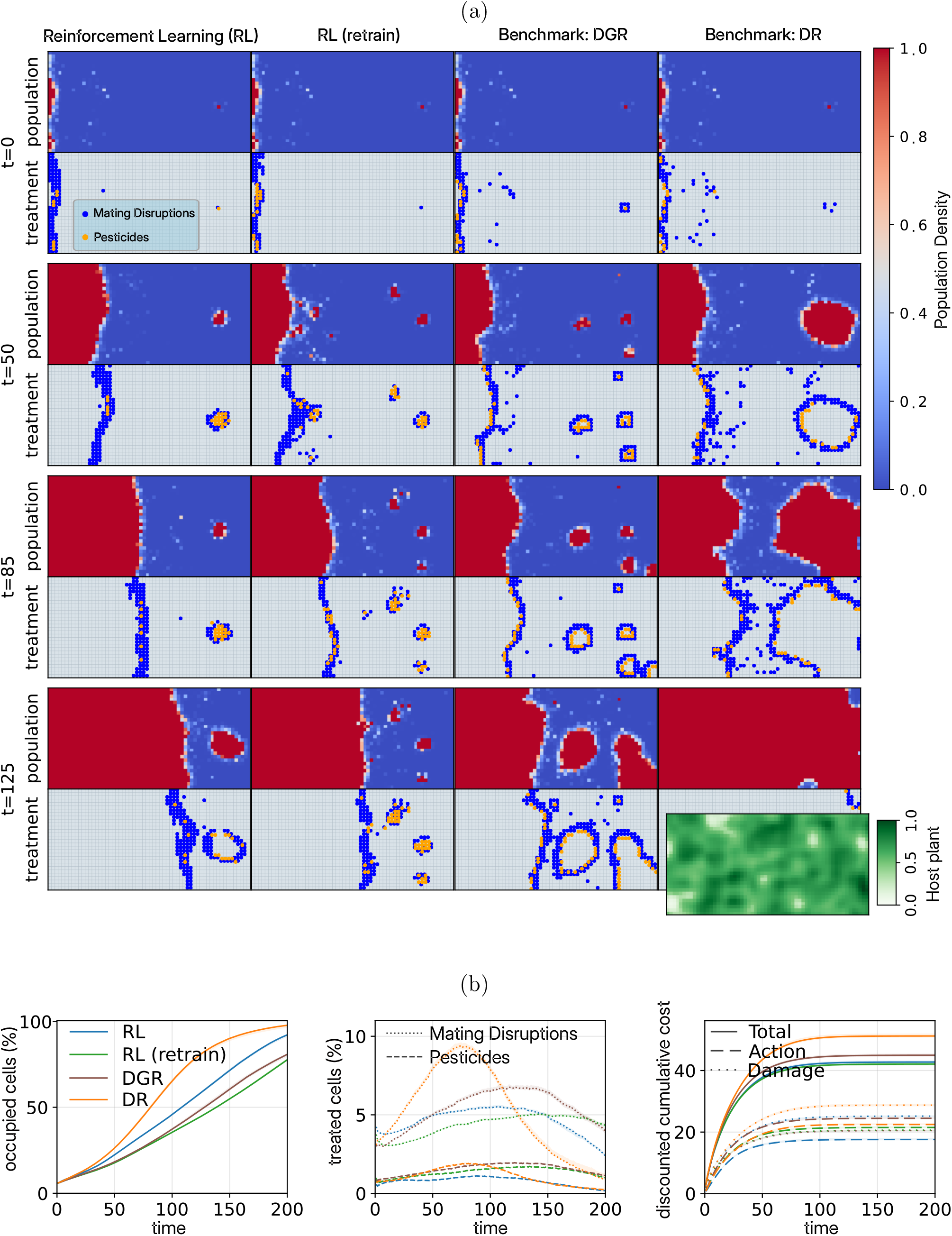
RL generalizes to large-grid tasks using broader spatial context. Representative 200-step evaluation on a heterogeneous-host 30 × 80 grid with stochastic off-front seeding. Columns compare a generalist RL agent and a retrained RL agent with two benchmarks based on local decision rules: DGR and DR. All simulations share the same initial state and host landscape. **(a)** Adult population densities and corresponding treatment actions are shown for the central 30 × 60 regions of the 30 × 80 simulation landscapes. Each column shows a simulation using a different policy, and each row corresponds to a different time step (*t* = 0, 50, 85, 125). The RL treatment maps show responses beyond fixed local density and gradient ranges. At *t* = 50 and *t* = 85, RL applies pesticides within high-density, low-gradient off-front colony interiors, but does not apply pesticides in main-front cells with similar local conditions. In contrast, the benchmarks apply fixed local rules that do not distinguish between the main front and off-front colonies. RL also applies pesticides along a line spanning a broad range of front densities (*t* = 85, 125) and reduces treatment near a low-host gap and on the colony side facing the main front (*t* = 125). The benchmarks instead produce repetitive boundary-following patterns, with more fragmented fronts and earlier outbreaks. **(b)** Demonstrated are several spatial aggregates averaged over 200 episodes for all four policies, including the percentage of occupied cells over time, the percentages of cells treated with mating disruption and pesticides, and discounted cumulative total, treatment, and damage costs. These averages compare mean policy performance across stochastic realizations, complementing the individual simulations in (a). Both RL agents achieve lower mean total costs than the two benchmarks. Relative to the generalist RL, the retrained RL’s policy spends more on treatment to reduce damage cost and improves spread suppression, yet its total cost improvement is minor.

### The RL policies combine front and off-front treatment strategies and outperform benchmarks

To assess the cost-effectiveness of the learned strategies relative to optimized local rules, we compare the generalist RL agent with benchmarks based on local population density (DR) or local density and its gradient over the eight nearest neighbors (DGR). Specifically, (i) the density-range benchmark policy (DR) applies treatment only if the population density in the cell lies within a certain range of values; (ii) the density-gradient-range benchmark policy (DGR) applies treatment only if both population density and its spatial gradient compared to its nearest neighbors lie within certain ranges. Note that, in both benchmarks, we consider one range for pesticide application and a distinct range for mating disruption. These ranges are optimized simultaneously via stochastic simulation, using the same objective function that was used to train the RL agent. To provide stronger benchmarks, we optimize them separately for each task, as explained in Methods. In contrast, the generalist RL agent is the same for all tasks. It receives no task label and selects actions solely from the fully observed state.

On the large-scale tasks, both range-based benchmarks reduce overall spread, but they cannot simultaneously slow the advancing front and contain off-front populations as efficiently as the generalist RL policy (Figure 2). The fixed treatment ranges needed for effective front control and off-front containment are incompatible. Choosing the benchmark treatment ranges therefore involves a trade-off: suppressing off-front populations necessitates targeting high-density cells, whereas treating such cells behind the invasion front is not cost-effective. Wider treatment ranges can capture these off-front populations, but also lead to overtreatment. Narrower ranges reduce this expense but may leave satellite populations insufficiently controlled.

The isolated colony management tasks (Figure 3, two rightmost columns) illustrate how fixed treatment ranges can lead to excessive treatment as colonies grow. Initially, all policies suppress nascent satellites and contain established colonies by targeting high-density cells.

**Figure 3.**
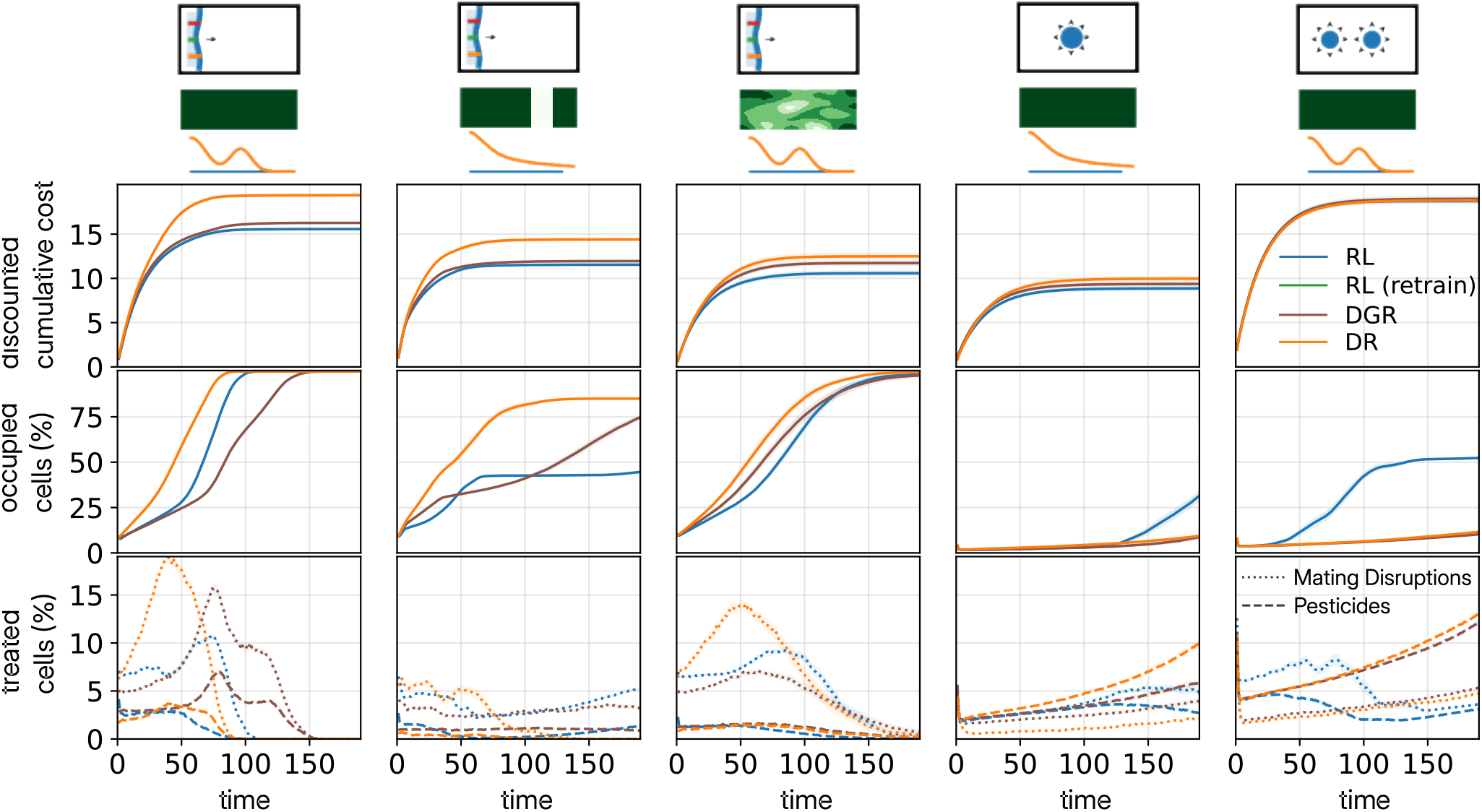
The figure compares RL and the benchmark policies (DGR and DR) on five invasion tasks. Rows 1–3 show discounted cumulative costs, percentages of occupied cells, and percentages of cells treated with mating disruption and pesticides, respectively. Results for each task are averaged over 50 simulations. The labels on the top identify the tasks and their dispersal kernels. We first consider the tasks in which invasion starts from the border (columns 1–3). When host-plant distribution is homogeneous (column 1), RL and DGR initially incur similar costs, but RL later achieves a cost reduction by relaxing treatment and allowing faster spread. When host-plant distribution is heterogeneous (column 3), RL gains an early cost advantage and slows the spread more effectively while using less treatment. When the host plants have a gap (column 2), the results shown are averages of left- and right-origin invasions, with the gap positioned toward the right. Both RLand DGR maintain treatment throughout the 200-year horizon. RL alone stops right-origin invasions, explaining why its mean occupied fraction remains near 50% when both directions are averaged. Yet its cost advantage over DGR remains small. In turn, in tasks where the invasive population begins from one or two colonies (columns 4 and 5), all policies begin with a short treatment pulse (see peak near time zero). Even under sustained containment, the colonies continue to grow, albeit very slowly, and maintaining containment requires treatment over an increasing area. The benchmarks progressively increase their treatment rates, whereas RL eventually withdraws.

As the colonies expand, DR and DGR treat an increasing area to maintain containment. In contrast, RL generally withdraws treatment earlier when larger or multiple initial colonies create a greater outbreak risk. Although the benchmarks’ containment strategies would not be cost-effective on large-grid tasks, the benchmarks remain comparably cost-effective in these training tasks because most of their additional treatment occurs late and is heavily discounted.

Host heterogeneity creates variation in spread rates and in the shape of the invasion front, increasing variation in density and gradients along population boundaries. Whereas RL responds to such variations, the benchmarks’ fixed treatment ranges must accommodate host-driven variation in abundance and gradients, creating a trade-off between treating important boundary cells and avoiding treatment of low-value cells. The first and third columns of Figure 3 illustrate the resulting advantage for RL. Under homogeneous hosts, RL and DGR initially have similar cost and spread trajectories, and RL gains a modest later advantage by relaxing control, allowing the front to advance faster. Under heterogeneous hosts, RL slows spread effectively and maintains a cost advantage throughout.

These differences in spatial control are reflected in the policies’ management costs on both training and generalization tasks (Figure 4). Across the training tasks, the generalist RL policy reduces mean cost by 16% relative to DR and 6.7% relative to DGR. For Generalization tasks, the corresponding reductions are 15% and 3.6%. Retraining RL provides a further improvement of two percentage points over the generalist policy. On late-detection generalization tasks, the generalist RL policy reduces mean cost by 20% relative to DR and 5.4% relative to DGR. Retraining improves performance by a further one percentage point, which suggests that the generalist RL policy has already learned how to combine front control with off-front management in these settings (Figure 2). Results for both long-distance kernels at 30% higher damage costs (Figure S3) show that RL’s cost advantage remains similar on training tasks but increases on larger-grid generalization tasks by 3.9 percentage points relative to DR and 2.3 percentage points relative to DGR.

**Figure 4.**
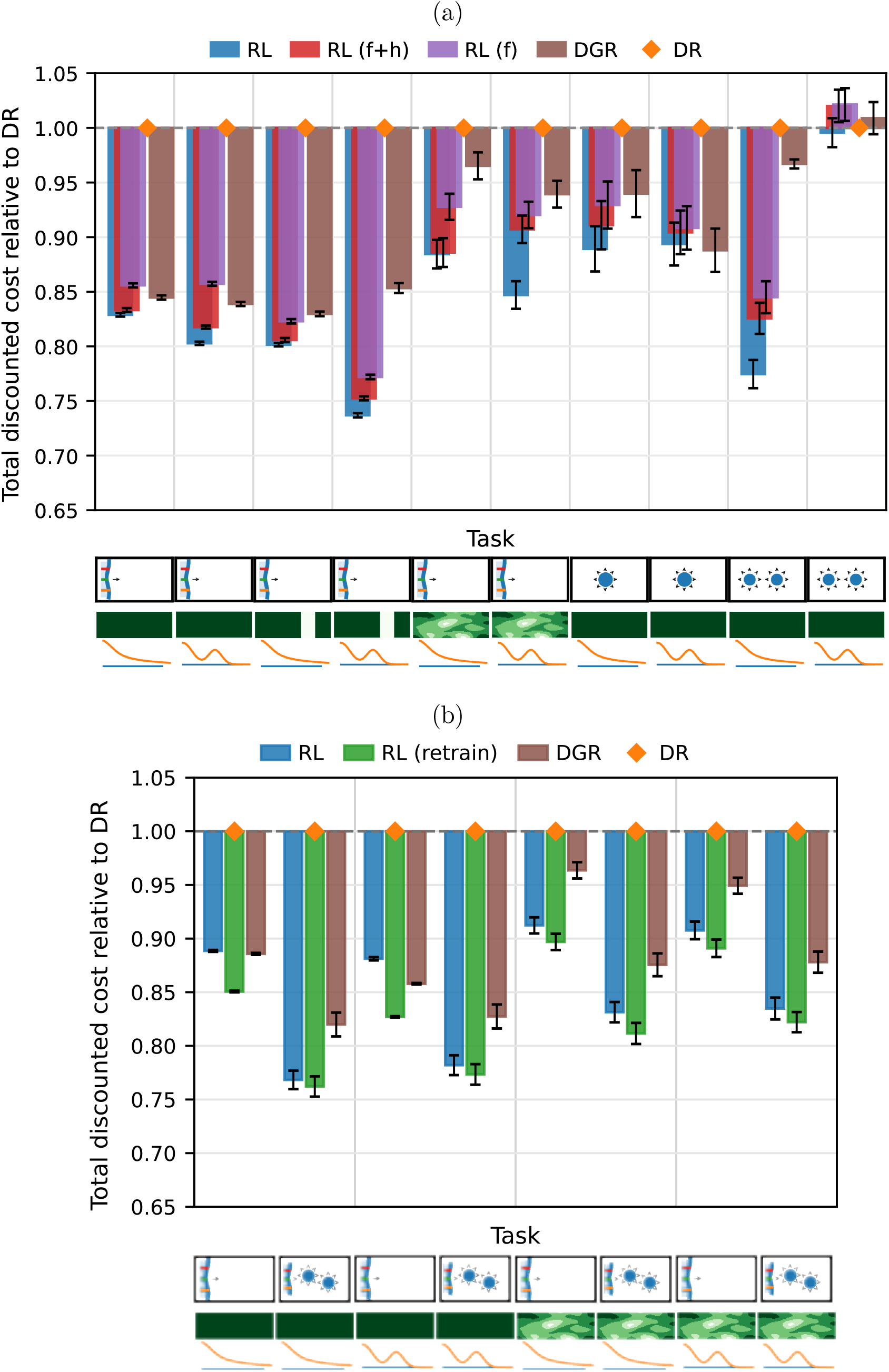
RL outperforms benchmarks in complex management settings. The discounted cumulative-cost ratios are shown relative to the DR benchmark. Lower values, indicating lower costs, are better. The bars are grouped by task. Each task label (bottom) specifies the invasion type (border invasion, single/double colony or border invasion + colonies), the host plant distribution, and the kernel used to model long-distance dispersal (Methods). Error bars show one standard error of each policy’s mean discounted cost across stochastic evaluation rollouts, normalized by the matched DR mean. **(a)** Training tasks on a 10 ×40 grid. The results are shown for two generalist RL agents, one for the fat-tailed kernel (tasks 1, 3, 5, 7, and 9) and one for the displaced Gaussian kernel (tasks 2, 4, 6, 8, and 10). Observation ablations test how performance depends on the available state information. The full RL policy observes adult density (*f*), invasion persistence (*i*), and host-plant density (*h*). Separately trained RL (f+h) and RL (f) generalist policies use the same task ensemble but observe only (*f, h*) and *f* , respectively. **(b)** Generalization tasks are evaluated on a 30 × 80 grid with invasions starting at the border, with or without seeding larger off-front populations. While generalist RL agents are evaluated directly on those tasks, RL retrain denotes the generalist RL agent retrained on each specific task (Methods). The generalist RL agents generalize successfully to the larger-grid tasks. Their cost ratios relative to DR are lower in settings with heterogeneous host distributions and additional off-front populations than in the uniform-host baseline.

### RL responds to an approaching front by redirecting treatment toward uninvaded areas

Our results show that RL employs long-range spatial coordination, adjusting treatment within off-front colonies in response to an approaching invasion front. Off-front colonies can accelerate range expansion. As the invasion front approaches, the colony’s front-facing side is likely to merge with the main front, whereas its away-facing side can still spread into unoccupied areas. Specifically, as the main front approaches, the front-facing side becomes less valuable to suppress because it is likely to be absorbed by the incoming invasion, whereas the away-facing side can seed further spread into unoccupied habitat. Therefore, treatment is more valuable on the away-facing side.

The RL policies correctly place more pesticides on the away-facing side than on the front-facing side near merging. In contrast, DGR places more pesticides on the front-facing side because higher-abundance cells occur more often there. Over the entire decade-long experiment, the away-facing edge advances by only one cell under RL policies. During the same decade, neighboring sections of the front catch up, contracting the merger bulge and straightening the front. These results demonstrate how RL learns to prioritize treatment according to a population’s contribution to future spread, rather than its current abundance alone.

## 3 Discussion

The principal finding is that RL can learn spatially contextual treatment strategies within a single policy that jointly manages the invasion front and off-front colonies. It identifies high-value local targets and adjusts treatment to each population’s changing contribution to future spread on a two-dimensional grid while responding to host heterogeneity. As the invasion progresses, it adjusts the timing, location, and extent of treatment, including limiting, redirecting, or withdrawing interventions to improve cost-efficiency (Figures 2 and 3). This flexibility is possible because the convolutional policy conditions each cell’s treatment on the observed state within a 17 × 17 spatial neighborhood. Unlike the benchmark policies, the neural network is not restricted to a single set of treatment ranges that must compromise between front and off-front control. Under the tested dispersal and invasion-damage cost assumptions, RL reduces mean total discounted costs relative to the optimized local benchmarks, particularly when front control and off-front colony management must be coordinated on heterogeneous-host landscapes (Figures 4 and S3).

The ecological principle reflected in the RL-trained policy is that the benefit of treating a location depends on the surrounding invasion and changes as the invasion progresses. Classical pest management relates treatment decisions to expected damage and control cost^54;55^. In a spreading population, expected damage also depends on how local populations contribute to future spread. Spatially explicit and network-based bioeconomic models formalize this dependence by accounting for how local control changes future spread, damage, and control costs elsewhere^17;18;24^ and reveal general principles of spatial invasion management: (i) management must anticipate consequences across both space and time; (ii) invasion geometry, particularly the exposed boundary, shapes control costs; (iii) the value of controlling one population depends on the surrounding invasion; and (iv) initial conditions and population dynamics determine the preferred management strategy. Extending models to large landscapes is computationally demanding, and even with simplified ecological dynamics, stochastic spread and control can substantially limit the landscape sizes tractable with exact optimization^24^. Our study demonstrates that RL can learn spatially contextual control policies at large spatial scales while incorporating complex population and treatment dynamics and management costs. Within a single simulation, we jointly recover the management principles identified by earlier bioeconomic models. Despite policy and value-function approximation, the RL policy proposes cell-level treatment decisions that generate consistent landscape-level responses reshaping the invasion front and populations. These responses generalize to the tested invasion configurations and larger landscapes, allowing the same learned RL policy to be applied without case-specific re-optimization (Figures 2, 4b and S3b). The policy does so without access to a critic, suggesting that it has learned transferable spatial treatment patterns.

The ability to coordinate different intervention types is central to sustainable pest management because it helps managers avoid unnecessary pesticide use while maintaining effective suppression^55^. The Slow the Spread program uses mating disruption as its main treatment, applying species-specific pheromones to selected low-density populations in the transition zone ahead of the generally infested area^40;48^. In our simulations, the agent mostly chooses no treatment at low abundances, uses mating disruption when mate limitation is effective, such as ahead of the invasion front or around intermediate-sized satellite colonies, and reserves pesticide for cases where direct suppression changes the future spread trajectory enough to justify its higher cost. The value of spatially separating these treatments was also demonstrated around host-plant gaps in a deterministic, one-dimensional model^45^. In our study, one example of context-dependent treatment is the change in treatment priorities within an off-front satellite colony as the invasion front approaches Figure 5. RL concentrates pesticide on the side contributing most to future expansion. It progressively reduces or withdraws pesticide from areas unlikely to contribute to future spread and manages them through mating disruption. A similar deprioritization based on expected contribution to future spread appears in the Slow the Spread Decision Algorithm, which assigns lower intervention priority to areas between the *action zone* and the *generally infested zone*^40^. The RL-trained policy therefore recovers a fine-scale version of this principle by allocating treatment type and intensity according to each population’s contribution to future spread rather than current abundance alone. Although our model is inspired by spongy moth dynamics, the management tactics we examine exploit the interplay between dispersal, host plants, and the Allee effect^42;56;57^. The resulting insights may therefore be relevant to other invasive species in which these mechanisms similarly shape establishment and spread.

**Figure 5.**
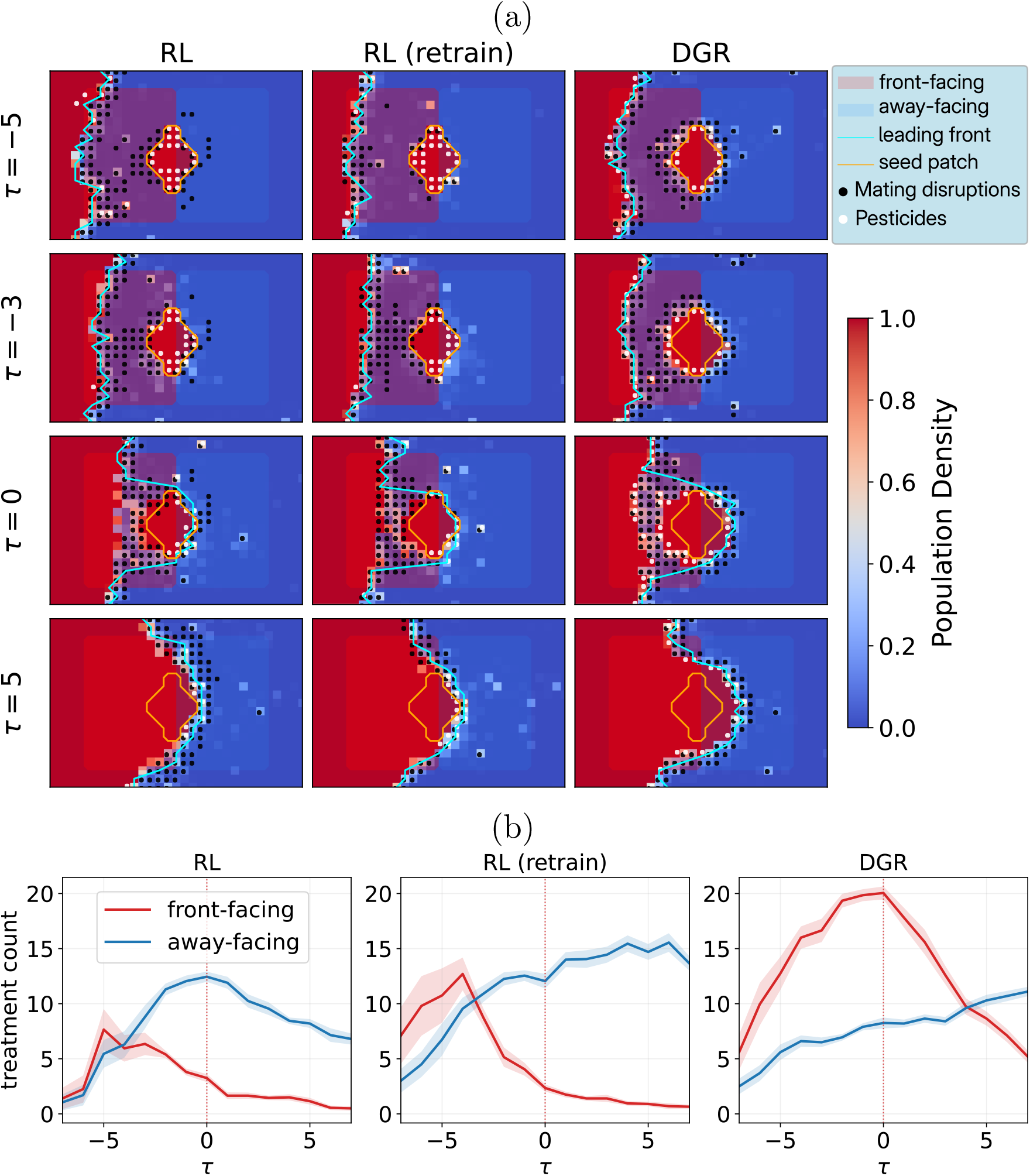
RL suppresses spread through long-range satellite–front coordination. A colony is placed ahead of an advancing invasion front and tracked as it merges with the front. Time *τ* is measured relative to merging (*τ* = 0). The figures compare the generalist RL agent trained only on the training tasks with the retrained RL agent and optimized DGR benchmark, for the task shown in Figure 2. **(a)** The figure shows adult density and the corresponding treatment locations under each policy. The shaded area around the introduced satellite colony is divided into two halves: the red region on the left faces the approaching invasion front (front-facing), and the blue region on the right faces away from the front (away-facing). Around the time of merging (*τ* ≈ 0), both RL policies concentrate pesticides on the away-facing region, whereas DGR concentrates pesticides on the front-facing region. Within the away-facing region, both RL policies preferentially apply pesticides to the most advanced parts of the front. Under both RL policies, the away-facing edge advances only one cell from *τ* = −5 to *τ* = 5. After merging, the front (turquoise line) straightens as lagging segments catch up. These patterns reveal long-range spatial coordination of the RL agent, which adjusts treatment within the satellite colony in response to an approaching invasion front. **(b)** Spatial aggregates averaged over 20 episodes show the numbers of pesticide-treated cells within the front- and away-facing regions as functions of the relative time *τ* , with bands showing one standard error. The red and blue curves correspond to the front- and away-facing regions, respectively. Consistent with (a), both RL policies increase treatment in the away-facing region before merging while markedly reducing treatment in the front-facing region from approximately *τ* = −4. By *τ* = 5, treatment is predominantly concentrated in the away-facing region. In contrast, DGR concentrates treatment on the front-facing region until after merging.

**Figure 6.**
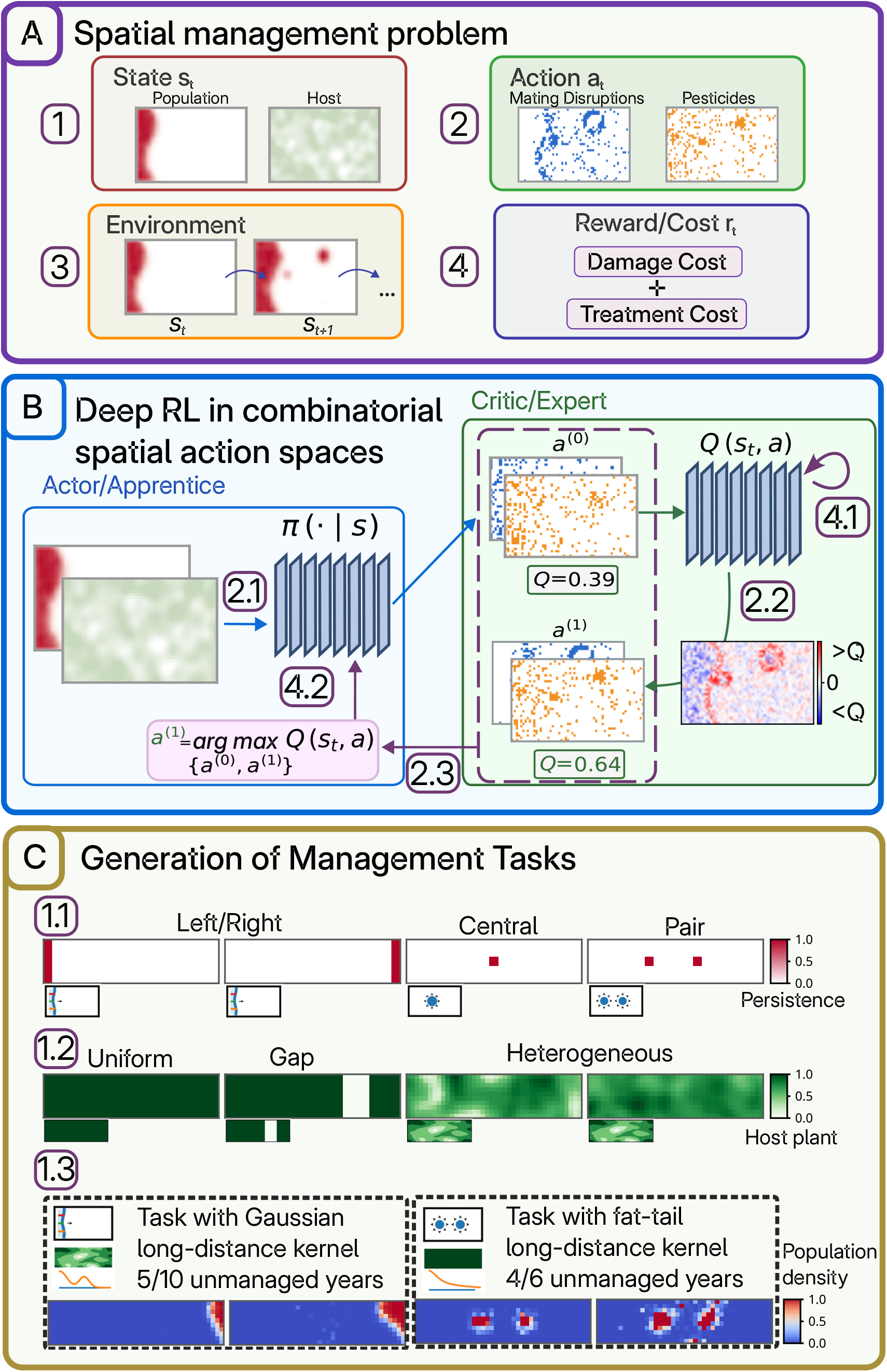
Spatial management, reinforcement learning, and task generation. (A) The spatial management control problem. The state *s*_t_ comprises adult density, invasion persistence (not shown), and host-plant density (1). The action *a*_t_ specifies where to apply mating disruption and pesticide (2). Population growth and dispersal under these treatments produce the next state *s*_t+1_ (3). The objective is to minimize expected discounted treatment and invasion-damage costs; the reward *r*_t_ is their negative annual sum (4). The numbering in (B) and (C) links task generation (1.1–1.3), action selection (2.1–2.3), and learning (4.1–4.2) to the state (1), action (2), and reward (4), respectively, rather than indicating chronological order. **(B) Action selection and learning**. The state enters the actor (apprentice) network *π* (stacked blue layers), which generates an initial action *a*^(0)^ (2.1). This action is passed to the critic (expert) network, which estimates its expected discounted return, *Q*. The gradient map indicates how local action changes affect *Q* (red, increase; blue, decrease), guiding the revised action *a*^(1)^ (2.2). The candidates in the dashed box are compared, and the action with the highest *Q* is selected (2.3) and fed back to train the actor by imitation (4.2). The critic’s separate learning loop updates its predictions using observed rewards and estimated future returns (4.1). **(C) Task generation**. Initial invasion-persistence patterns seed adults (1.1) on host-plant landscapes (1.2). An unmanaged period of growth and dispersal produces the adult-density fields at the start of management (1.3). Task labels identify the invasion pattern, host landscape, and long-distance dispersal kernel.

Similar large landscape-scale applications in conservation and forest management demonstrate the relevance of AI’s role beyond invasive species. For example, the CAPTAIN project uses evolutionary policy search to learn conservation or restoration priorities^34;58^. Lauer et al.^59^, in turn, optimize timber-harvesting and fuel-treatment decisions across a fire-threatened forest landscape using reinforcement learning that combines a spatially factorized post-decision value approximation with hybrid simulated-annealing/tabu search. Both approaches share model parameters across spatial units, with spatial context entering through predefined aggregation functions. In CAPTAIN, a shared multilayer perceptron learns a nonlinear mapping from ecological features to planning-unit priorities. Lauer et al.^59^, by contrast, approximate post-decision values as a weighted sum of predefined linear or nonlinear basis functions. Our framework likewise shares model parameters across space but aims to reduce reliance on problem-specific structure, including predefined aggregation functions, explicit post-decision decompositions of management effects, and prescribed value-function bases. To this end, we use deep convolutional networks to learn spatial features and state–action values from simulator experience. This flexibility, supported by approximately half a million trainable parameters per agent, comes at the cost of computationally demanding training. However, remaining agnostic about which spatial relationships should govern treatment priorities allows the framework to learn coordination patterns that are not prescribed in advance, as illustrated by the long-range spatial coordination observed in our simulations.

The framework developed here should be interpreted as decision support rather than field-ready automation. Real-world reinforcement learning faces challenges involving distribution shift, partial observability, safety constraints, reward specification, and policy validation, with potentially costly mistakes and slow feedback in ecological management^50;60^. Policies learned under specified assumptions about population dynamics, treatment effects, and management costs may exploit simulator simplifications, and their effectiveness may decline when ecological or operational conditions differ from those represented during training. Operational use also requires accounting for imperfect surveys and delayed detections rather than assuming full-state information. This motivates an extension to policies defined over belief states, with monitoring and treatment decisions optimized jointly. The policy could then balance the cost of monitoring against its expected benefit for subsequent management outcomes^61^.

Our results provide a proof of concept for RL as an optimization framework for large-scale spatial management. Grounding both ecological dynamics and the observation process in existing trapping records would provide a basis for developing and empirically testing landscape-scale management strategies. More broadly, the framework could be adapted to other invasive species and to spatial resource-management problems in which local actions have consequences elsewhere. Its potential lies in coordinating interventions across space and time to achieve greater long-term benefits from limited management resources.

## 4 Methods

### 4.1 Model dynamics

In this section, we provide a complete description of the model. Our spongy moth dynamic model incorporates local reproduction subject to a strong Allee effect due to mate-finding failure, and stratified dispersal, in which stochastically generated colonies grow and lead the invasion front.

#### 4.1.1 Discrete-time dynamic equations

We assume that the population is characterized by an adult population density, without modeling all life stages of the species or differentiating between males and females. This is a common practice for modeling univoltine insects, as each life stage occurs in a different season, and treatment at any stage is ultimately translated into changes in adult density^62^. Specifically, the model describes, in each grid cell *i, j* and at time *t* (years), the adult density 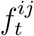, the host-plant density *h*^ij^, which we assume to be constant over time, and the invasion persistence 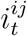, which characterizes a hidden population that does not respond to treatment and provides a second source of adults. In particular, in some invasive insects, individuals can persist in inaccessible refuges that escape effective pesticide treatment, allowing populations to recover after control efforts^63;64^. This is one reason why eradication is generally more likely to succeed when invasions are detected early and remain spatially restricted^65;66^. In our model, we incorporate this effect to represent populations that can recover after treatment, so that local suppression need not imply permanent eradication. This allows us to study sustained management on finite landscapes, including populations away from the main invasion front.

The persistence variable also determines delayed invasion-damage costs (Section 4.1.5) and is used to initialize populations (Section 4.2.4).

We represent the fields in the state vector ***s***_t_ = (*i*_t_, *f*_t_, *h*)^⊤^ ∈ *S*. The dynamics are described through three stages applied to the state vector: local reproduction, denoted by the operator *ℛ* (Section 4.1.2); coupled local persistence-growth and seeding, denoted by *P* (Section 4.1.2); and dispersal, denoted by *K* (Section 4.1.3). With this, the update equation becomes

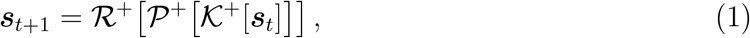

i.e., we first let the species disperse, then update invasion persistence and seed additional adults, and finally apply reproduction. The superscript + denotes carrying capacity adjustment at each stage of the dynamics. Specifically, for a single stage *ℱ* = *ℛ, P, K* and a normalized density component *x* ∈ {*i, f* }, the operator + imposes logistic-type saturation by reducing population density increments near carrying capacity, while leaving reductions unchanged:

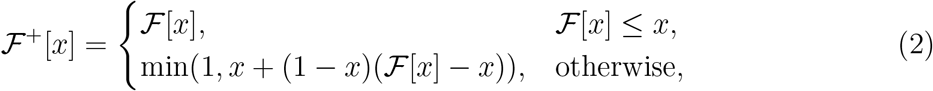

where we use the shorthand *ℱ*[*x*] = (*ℱ*[***s***])_x_.

#### 4.1.2 Local population growth

Local population growth combines density-dependent mating with host-dependent reproduction. Given the local adult density *f* ^ij^, the reproduction operator *ℛ* is defined through

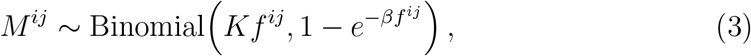

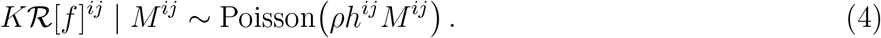

Here, *K* is the carrying capacity, which is assumed constant, and *M* ^ij^ is the sampled number of mated females from *Kf* ^ij^ trials, one per adult, with mating success probability 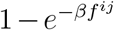 in each trial. *Kℛ*[*f*]^ij^ is the sampled number of offspring generated by the mated females, and *ℛ*[*f*]^ij^ is its associated density, obtained by dividing the sampled number by *K*. Only then do we apply the carrying-capacity adjustment in Equation (2) to ensure that the resulting density *ℛ*^+^[*f*]^ij^ ∈ [0, 1].

The coefficient *β* controls the density dependence of mating; accordingly, the term 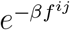 characterizes the probability that a female is not found by any male ^62^. At large *f* ^ij^, this probability approaches zero, making mate-finding limitation negligible. Each mated adult produces *ρh*^ij^ offspring on average, with *ρ* giving the expected number at full host density. We note that adults are removed after reproducing, meaning their offspring become the adults of the following year. We do not explicitly model mortality other than through Equation (2).

Adult and host-plant densities drive growth in invasion persistence, which in turn seeds additional adults. This process is defined through the operator *P* as follows:

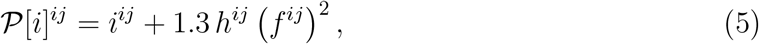

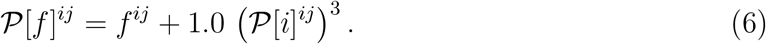

These exponents and coefficients were chosen to make persistence accumulate gradually and to keep persistence-driven seeding of adults small until *i*^ij^ ≈ 1.

#### 4.1.3 Dispersal components

Dispersal redistributes adult density across the grid through long- and short-distance movement. We model these movements as two stages using the same stochastic construction, in which the density leaving each source cell is sampled and redistributed among destination cells.

For a pre-stage adult-density field *f* , we sample the outflow from cell (*k, l*) according to

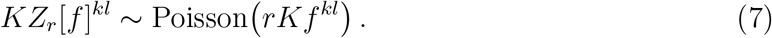

Here, *KZ*_r_[*f*]^kl^ is the sampled number of adults dispersing from cell (*k, l*), and *Z*_r_[*f*]^kl^ is its associated density, obtained by dividing the sampled number by *K*. The parameter *r* denotes the dispersal rate, so the expected outgoing density is *rf*^kl^.

The number of destination draws *L* is sampled independently for each source cell (*k, l*) in long-distance dispersal and fixed in short-distance dispersal (Table 1). The destination coordinates are distributed as

**Table 1:** Population-dynamics, dispersal, and management parameter values. Kernel distance parameters and the hard cutoff *R* are measured in grid-cell units. Short-distance settings are shared by both long-distance variants. Costs refer to the baseline setting.

| Parameter | Symbol | Description | Values |
| --- | --- | --- | --- |
| *Local population dynamical rates* |  |  |  |
| Adult carrying capacity | $K$ | Maximum number of adults | $7.5 \times 10^6$ |
| Mate finding | $\beta$ | Density-dependence coefficient | $7.5 \times 10^{-4}$ |
| Reproduction | $\rho$ | Offspring per mated adult at full host density | $10^4$ |
| *Dispersal* |  |  |  |
| Long distance, fat-tailed | $a, b, s, R, r, L$ | Distance scale; tail decay; radial offset; cutoff; dispersal rate; destination draws | 10, 2.5, 0, 30, 0.60, Poisson(0.6) |
| Long distance, Gaussian | $\sigma, s, R, r, L$ | Kernel width; radial offset; cutoff; dispersal rate; destination draws | 1.7, 16, 30, 0.43, Poisson(0.6) |
| Short distance, Gaussian | $\sigma, s, R, r, L$ | Kernel width; radial offset; cutoff; dispersal rate; destination draws | 2, 0, 2, 1.00, 30 (fixed) |
| *Management objective* |  |  |  |
| Pesticide cost | $c_p$ | Cost per cell | 0.045 |
| Mating disruption cost | $c_m$ | Cost per cell | 0.00089 |
| Invasion damage cost | $c_d$ | Damage per cell when persistence crosses 0.8 from below | 0.11 |
| Discounting | $\gamma$ | Discount factor | 0.95 |

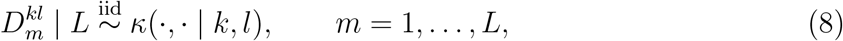

where *m* indexes the destination draws from source cell (*k, l*) and *κ* is the stage-dependent dispersal kernel (Section 4.1.4).

For *L >* 0, each draw assigns an equal share 1*/L* of the outgoing density to its sampled destination. A destination cell may be sampled more than once. Thus, the fraction of the outgoing density sent from (*k, l*) to (*i, j*) is given by:

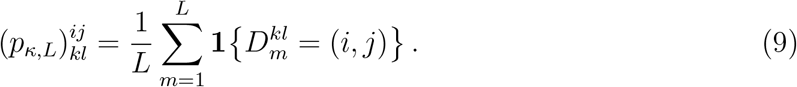

This redistribution occurs simultaneously across the grid, so a cell can both send adults to other cells and receive adults from multiple sources. The dispersal operator *K*_(r,κ,L)_ gives the adult-density field after this redistribution:

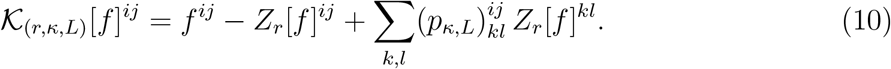

The full dispersal operator applies long-distance dispersal first, followed by short-distance dispersal,

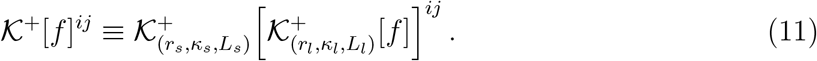

Note that satellite colonies are primarily produced by the 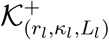 operator, whereas a potential invasion front would move via the 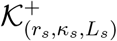 operator, accelerating as it merges with these colonies.

The resulting spread is illustrated in Figure S1 under uniform full host coverage (*h*^ij^ = 1.0), which gives the fastest expected spread for a given kernel and treatment regime. Each simulation step represents one year, and assuming a cell size of approximately 12 km × 12 km gives spread velocities comparable in magnitude to historical unmanaged estimates and managed rates in the northern region of the Slow the Spread program (see Figure S1).

#### 4.1.4 Dispersal kernels

We model long-distance dispersal using two alternative kernels: Gaussian and fat-tailed. Short-distance dispersal is modeled using a Gaussian kernel. The corresponding base kernels, denoted by 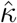, are given by

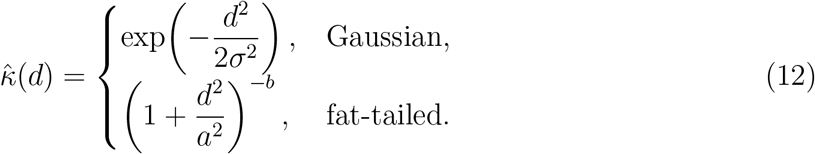

Here, *d* denotes Euclidean distance in grid-cell units, while *σ, a*, and *b* are kernel parameters whose descriptions and values are given in Table 1.

Motivated by the Coalescing Colony Model^8^, we introduce a radial offset *s* in the Gaussian long-distance kernel. The subsequent short-distance stage redistributes both adults remaining near the source and those transported over longer distances. Together, these stages produce an effective dispersal profile with a near-source peak and a second peak at a distance approximately equal to *s*. The latter promotes the formation of satellite colonies away from the main population. For the fat-tailed kernel, we set *s* = 0.

The finite grid introduces boundary effects, which we account for when constructing each dispersal kernel *κ* from its base kernel 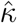. On the one hand, discarding outflow outside the grid makes the boundary an artificial population sink. On the other hand, renormalizing the kernel over in-grid destinations can bias movement toward the interior. To mitigate these artifacts, we retain all outflow within the grid and apply a degree correction to reduce inward bias:

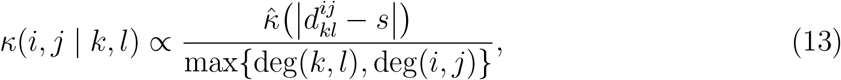

where 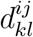 is the Euclidean distance from source cell (*k, l*) to destination cell (*i, j*). Destinations are restricted to those within a hard cutoff distance *R*, and deg(*x, y*) is the number of admissible destinations from cell (*x, y*) within this cutoff.

#### 4.1.5 Interventions and management objective

The two available interventions differ in terms of their mechanism, timing and cost. Pesticide application targets the larval stage, which we model by reducing the density of adults by 80% in treated cells before the reproduction stage (Equation (1)). Mating disruption involves deploying pheromone dispensers to confuse male moths. We model this effect by replacing *β* with 0.75*β* in Equation (3)^62;67^ in treated cells for one year, thereby reducing the probability of mating during the reproduction stage. Both interventions can be applied to the same cell simultaneously. Mating disruption has a low cost per cell, enabling its widespread application. Pesticide application costs approximately 50 times as much per treated cell, making selective spatial targeting particularly important for cost-effective use.

The annual reward is the negative of the total treatment and invasion-damage costs:

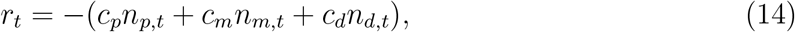

where *n*_p,t_ and *n*_m,t_ count cells receiving pesticide and mating disruption in year *t*, respectively, and *n*_d,t_ counts cells whose invasion persistence crosses 0.8 from below during year *t*. The coefficients *c*_p_ and *c*_m_ specify the per-cell treatment costs, and *c*_d_ specifies the invasion-damage cost (Table 1).

The objective is to maximize the expected discounted return:

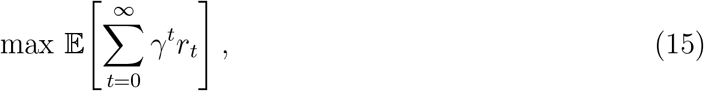

where *t* = 0 marks the start of management, and *γ* is the discount factor.

### 4.2 Reinforcement learning

#### 4.2.1 RL for large spatial control problems

We optimize the spatial control problem using a reinforcement-learning algorithm that combines ideas from learning in large discrete action spaces^38^ and search-based policy iteration^52;53^. Our approach builds on off-policy, value-based reinforcement learning^27;32^. The algorithm uses two learned networks, an actor and a critic. The actor *π* proposes actions, while the critic *Q* estimates the expected discounted return given a state and action. The learning procedure involves experience collection (Figure 6 **A**), during which transitions generated through interaction with the simulator are stored in a replay buffer, and training (Figure 6 **B**), during which transitions are sampled from that buffer and used to update both networks. We additionally use prioritized replay^68^ and maintain slowly updated target copies of both networks^69^.

The challenge lies in selecting landscape-wide actions from a discrete action space whose size grows exponentially with the number of grid cells, making exhaustive evaluation infeasible for large landscapes. For a large but fixed discrete set of actions, Dulac-Arnold et al.^38^ propose an approach in which an actor trained using deep deterministic policy gradient generates a proposal in a continuous embedding of the action set. The embedding defines neighborhoods among actions, allowing a nearest-neighbor search to retrieve a small set of discrete candidates. A critic then selects the highest-valued candidate for execution. Alternatively, the actor can be trained as an apprentice to imitate actions selected by a search-defined expert^52;53^. The resulting feedback loop allows the actor to learn from the more costly search and, in turn, provide better proposals for subsequent searches.

We design such a critic-guided search that starts from the actor proposal and operates directly on discrete spatial maps. We train the actor by imitation to reproduce the selected discrete action. For computational efficiency, the actor selects the full landscape-wide action per year in a single neural forward pass. The procedure remains computationally tractable and applicable as landscape size and action dimension grow. The local search implementing this procedure is described next.

Let *a* ∈ {0, 1, 2, 3}^XY^ denote an action on an *X* × *Y* grid, where the four values encode the possible combinations of the two interventions in each cell. The actor serves as an *apprentice* and samples an initial action map

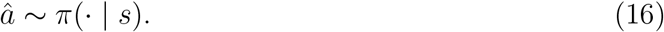

In our implementation, *π* is a factorized Bernoulli proposal distribution with one logit *ℓ*_k_(*s*) for each action bit *k* and corresponding probability Pr(*â*_k_ = 1 | *s*) = *σ*(*ℓ*_k_(*s*)). We use the critic as an *expert* by acting approximately greedily with respect to *Q*, that is, seeking the action with the highest predicted return starting from *s*. Rather than evaluating all 4^XY^ possible maps, the search starts from Equation (16) and explores a limited candidate set C(*s, â*).

The search generates new candidate actions through bit flips using gradients of the regularized surrogate *J*(*s, a*) = *Q*(*s, a*) + *λ* log *π*(*a* | *s*). A bit flip switches one intervention on or off in a single grid cell. The critic term *Q*(*s, a*) favors actions with higher predicted returns, whereas the *λ* term encourages learning to proceed through smaller changes from the current proposal distribution (16). Each search changes at most *n*_flip_ = 6 treatment decisions. The resulting expert action provides the imitation target *a*^⋆^ = arg max_a∈C(s,â)_ *Q*(*s, a*), which the apprentice is trained to imitate via the loss ^52^:

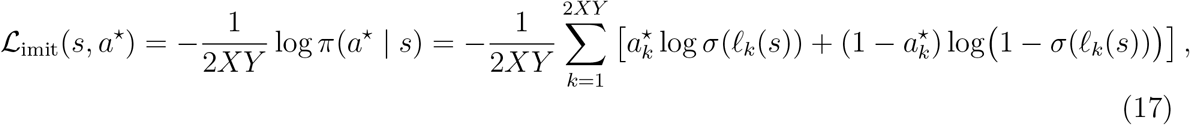

where the right-hand side is the mean binary cross-entropy between the actor’s predicted action probabilities and the expert’s action bits.

The procedure is organized in search rounds, each refining action candidates from the previous round through further bit flips. We use one search round during experience collection and two rounds to select the next actions used in the critic’s bootstrap targets during training. Furthermore, over the course of training, *λ* is gradually increased from 0 to 0.02. Convergence results in an apprentice that provides effective management actions directly, without the need to invoke either the critic or the search.

#### 4.2.2 Network architecture

The actor and critic neural networks are fully convolutional and contain no dense layers. The actor maps the ecological state *s* to a logit tensor *ℓ*(*s*) ∈ R^2×X×Y^ , with one channel for mating disruption and one for pesticide application. The critic receives *s* and *a* and produces a map *q*(*s, a*) ∈ R^X×Y^ , whose entries are summed over cells to obtain 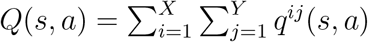.

Both networks use four residual blocks^70^. Each block contains two 3 × 3 convolutional layers with 64 hidden channels and ReLU activations. Including the output layer, each per-cell output has a receptive field of 17 × 17 cells, meaning that each output cell depends on state and action information within a neighborhood of this size. Because both networks are fully convolutional, the same weights can be used with different grid sizes and hence different action dimensions. The spatial context available to each per-cell output nevertheless remains limited to the fixed receptive field.

#### 4.2.3 Training setup and parameters

Each generalist agent completed 9,000 rollout sessions. Each session used 16 parallel workers, each of which collected 150 steps, yielding approximately 22 million transitions per agent. For comparison, the RL agents are retrained on the generalization tasks collecting approximately 7 million additional environment steps. We use commonly adopted replay, loss, optimization, target-network, and exploration hyperparameters^32;69^, as summarized in Table 2. Learning curves for the different generalist agents are given in Figure S4.

**Table 2:** Training hyperparameters.

| Hyperparameter | Value |
| --- | --- |
| Replay capacity | $10^6$ transitions |
| Replay warmup | 20,000 transitions |
| Batch size | 32 |
| Prioritized replay | $\alpha_{\text{PER}} = 0.3, \beta_{\text{IS}} = 1$ |
| Huber loss | $\delta = 1$ |
| Gradient clipping | norm 10 |
| Actor learning rate | $6 \times 10^{-5} \rightarrow 10^{-5} \rightarrow 3 \times 10^{-6}$ |
| Critic learning rate | $10^{-4} \rightarrow 5 \times 10^{-5}$ |
| Polyak coefficient | $0.005 \rightarrow 0.002 \rightarrow 0.001$ |
| Exploration | $\epsilon : 0.8 \rightarrow 0.005$ |

#### 4.2.4 Task generation

Training episodes are drawn from five procedurally generated tasks (Figure 6 **C**). Each task involves an invasion pattern combined with a distribution of host plants on a rectangular grid measuring 10 × 40. We distinguish between two types of initial invasion pattern: border invasions and isolated colonies.

A border invasion describes a population that initially covers most of the grid’s height on either the left-hand side or right-hand side and spreads horizontally afterwards. An isolated colony initially occupies four or more connected cells, but does not span the full height or width of the grid. It spreads in all directions. The first three tasks combine a border invasion with (i) uniform host coverage; (ii) uniform host coverage interrupted by a host gap near the right edge; (iii) spatially heterogeneous host coverage comprising sparse and dense areas. The other two tasks use uniform host landscapes and involve (iv) one isolated colony in the central area, and (v) two nearby but separate colonies in the central area. We note that more task combinations can be formed. Thus, the generalist RL agent is not trained on all possible combinations; for example, it is not trained on isolated colonies in heterogeneous-host landscapes.

Each episode begins with an unmanaged warmup period, during which the population grows and spreads stochastically without intervention. As a result, repeated episodes generally begin management from different states and may include satellite colonies generated by long-distance dispersal. Within each task, the warmup duration and, for border invasions, the invasion side (left or right) define the task variants. We use 4 and 6 warmup steps for all tasks except the heterogeneous-host task, which uses 5 and 10 because spread and population buildup are slower. Thus, each border-invasion task has four variants and each colony task has two, giving 16 variants in total.

Generalization tasks comprise four tasks on a 30 × 80 grid. Using the same border-invasion and host-landscape generators as in training, they combine uniform or heterogeneous host coverage with the absence or presence of stochastic off-front seeding. Each task has left- and right-origin variants and uses one warmup duration, 4 steps for uniform hosts and 5 for heterogeneous hosts. At each time step in a seeded task, an auxiliary population is initialized at carrying capacity in each cell (*i, j*) with probability 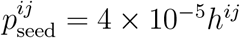. This population then spreads for three consecutive time steps using the short-distance operator 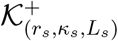 in (11) before being converted into invasion persistence, which continually seeds adults.

To examine policy learning under selected alternative assumptions about dispersal dynamics and damage costs, we consider two long-distance dispersal kernels (fat-tailed and Gaussian) and two damage-cost levels. The baseline cost setting, defined in Section 4.1.5, is used in the main text, while results for a 30% increase in damage costs are provided in Figure S3. For each of the four kernel–cost combinations, we trained three generalist RL agents: one full-observation agent and two agents with observation-ablated actors. All agents were trained on the same 16 task variants and evaluated under the kernel and damage-cost settings used for their training. The observation ablations in Figure 4a assess sensitivity to the state variables available to the RL agent. During evaluation, the agent selects actions directly, without the critic or search. The full-observation agent receives adult density (*f*), invasion persistence (*i*), and host-plant density (*h*). The RL (f+h) agent receives (*f, h*), whereas the RL (f) agent receives only *f* . In all cases, the expert retained access to the full state (*i, f, h*) during training.

#### 4.2.5 Benchmark policies

The DR and DGR benchmarks use the same treatment-specific ranges in every cell. Both policies make separate decisions regarding mating disruption and pesticide, allowing either treatment, both, or neither. The DR policy has four parameters, corresponding to lower and upper local adult-density bounds for each treatment. DGR adds a treatment-specific range for local density-gradient magnitude. Each treatment therefore has four parameters—two density bounds and two gradient bounds—giving eight in total. DR applies a treatment when local adult density lies within its optimized range, whereas DGR applies it only when both density and gradient magnitude lie within their respective ranges. The gradient feature is computed as the average absolute density difference between the target cell and its eight surrounding cells.

For each training and large-scale task variant, the parameters of each benchmark are optimized separately using the objective in (15) and ASTRO-DF^71;72^, as implemented in SimOpt^73^. We run each optimization 5–10 times with different random seeds and retain the solution with the best objective value. We verified that runs with different seeds yield policies with comparable final performance. As an optimizer cross-check, we verified the selected training-task optimizations using Dragonfly ^74^.

The task-level costs reported in Figure 4 are unweighted means across task variants, combining warmup durations and, where applicable, invasion sides. Although the benchmark rules are left–right symmetric by construction, fitting them separately allows their optimized ranges to differ between invasion sides. The generalist RL policy instead uses one policy across variants and has no built-in left–right symmetry, so it must learn how to respond to both invasion sides. The host-gap task itself is asymmetric because the gap remains near the right edge, resulting in different benchmark ranges, cost and spread trajectories (Figure 3).

## Data availability

This study used computational simulations. Simulation parameters are provided in Methods. The simulation environment and trained agent weights needed to regenerate the principal results will be made available as described in Code availability.

## Code availability

The ecological simulation code, the trained reinforcement-learning policy used for the analyses, and scripts for loading the policy will be made available on Zenodo after publication.

## Author contributions

J.-N.L. and A.L. conceived the study. J.-N.L. developed the software, performed the simulations, and analysed the results. A.L. supervised the study. J.-N.L. and A.L. contributed to writing and revising the manuscript.

## Ethics and consent

Ethics, Consent to Participate, and Consent to Publish declarations: not applicable.

## Supplementary figures

**Figure S1:**
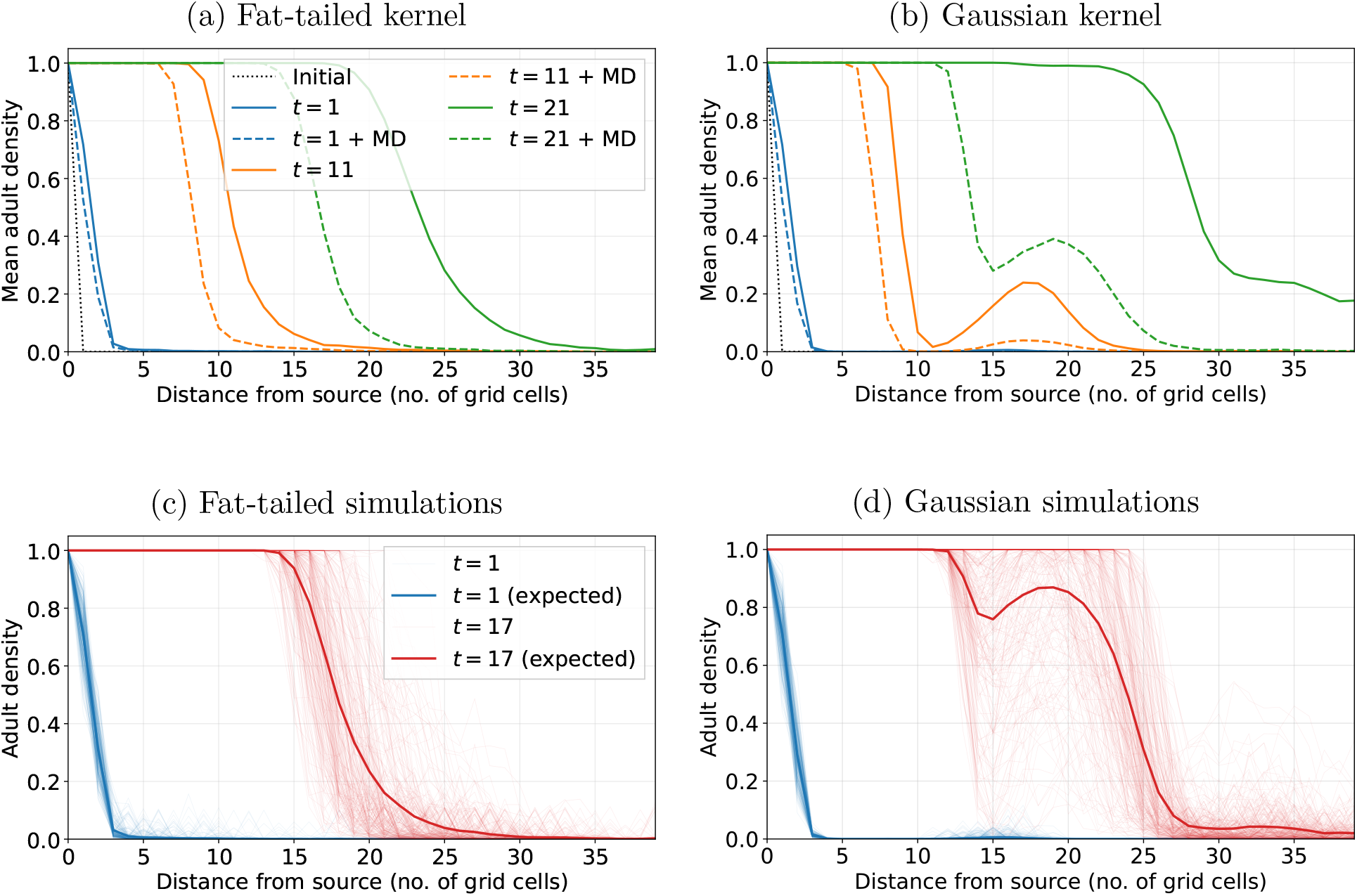
Spread under fat-tailed and Gaussian dispersal from a localized initial population at distance zero on the 10 × 40 grid. Host-plant density is uniform at *h*^ij^ = 1.0, enabling the fastest expected spread for each dispersal kernel and treatment regime. Each simulation step represents one year, and each grid cell is assumed to be approximately 12 km × 12 km. **Top row:** Expected adult density at years 1, 11, and 21 without intervention (solid lines) and with intensive mating disruption (dashed lines). **Bottom row:** Individual simulations at years 1 and 17 without intervention; thicker lines show expected profiles. Mating disruption slows spread under both kernels but does not stop it. The fat-tailed kernel initially spreads faster but produces diffuse, low-density satellites. The Gaussian kernel produces larger satellites that are more likely to persist and later merge with the main front, thereby accelerating the spread more than the fat-tailed kernel. Tracking the 0.5-density crossing of the expected profiles between years 11 and 21 gives front advances of approximately 12.4–19.7 grid cells without intervention and 6.4–8.4 grid cells under mating disruption, corresponding to mean velocities of approximately 15–24 and 8–10 km yr^−1^, respectively. The untreated velocities are comparable in magnitude to the historical reference rate of 19.6 km yr^−1^ reported by Coleman et al.^48^ (see also Tobin^49^), while those under mating disruption are comparable to the northern-region mean of 9.62 km yr^−1^ during 2000–2020.

**Figure S2:**
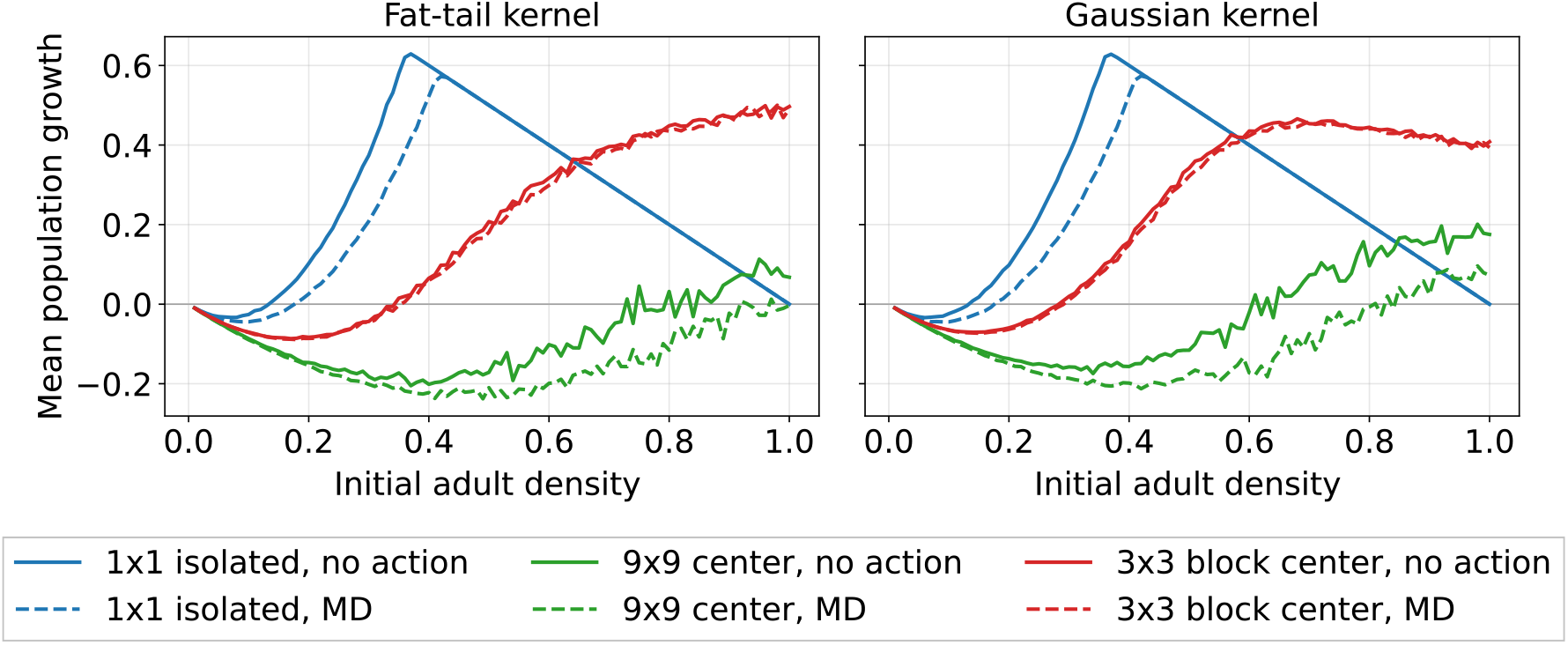
Mean single-step population growth as a function of initial adult density under fat-tailed (left) and Gaussian (right) dispersal. Colors distinguish an isolated 1 × 1 cell, a single occupied center cell on a 9 ×9 grid, and the center of an occupied 3 ×3 block. Solid lines show no intervention; dashed lines show mating disruption. Dispersal into empty surroundings reduces expected local growth relative to the isolated 1 × 1 cell, whereas immigration from occupied neighbors partly offsets this loss. Mating disruption (MD) further suppresses growth, with a smaller effect at the center of the occupied 3 × 3 block.

**Figure S3:**
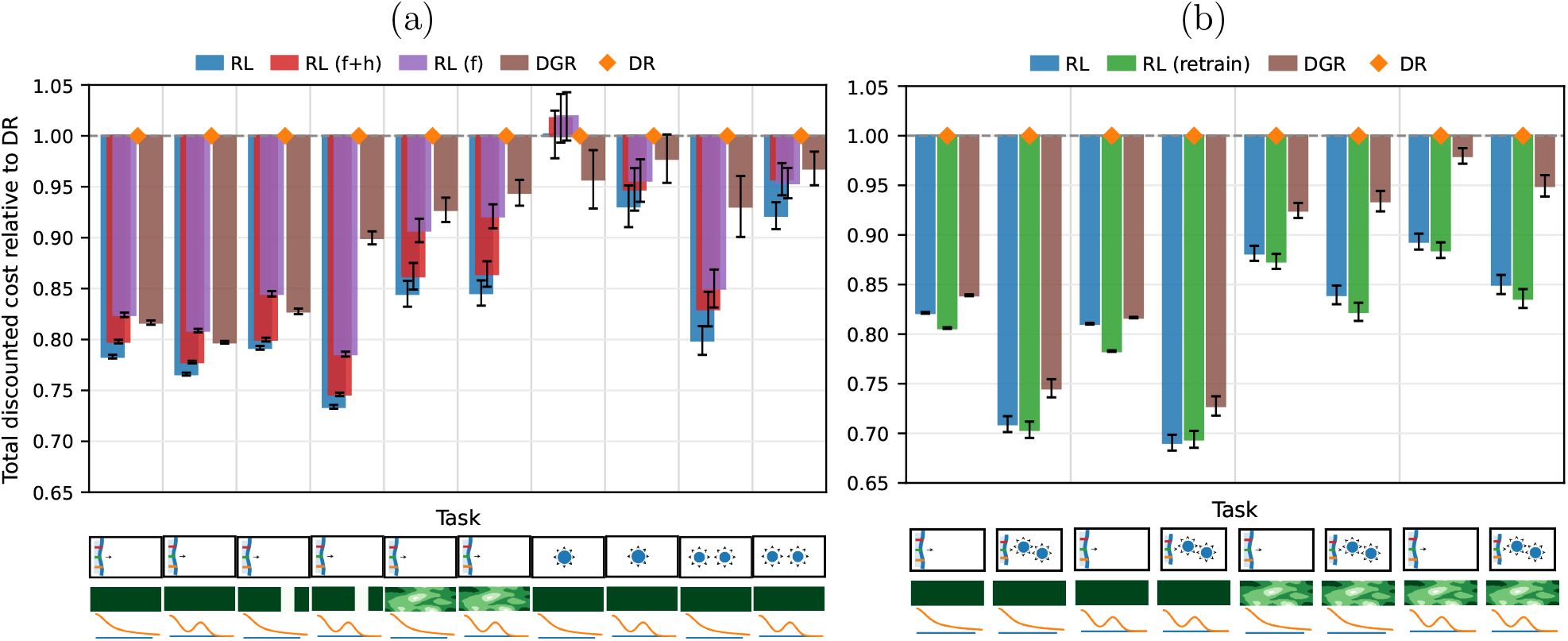
RL outperforms benchmarks in complex management settings under 30% higher damage costs. Discounted cumulative-cost ratios relative to the matched DR benchmark for **(a)** training tasks on a 10 × 40 grid and **(b)** generalization tasks on a 30 × 80 grid. Lower values indicate lower costs; error bars show one standard error. Task labels and policy definitions follow Figure 4. The generalist RL policies achieve mean cost reductions relative to DR and DGR, respectively, of 16% and 6.9% on training tasks and 19% and 5.9% on generalization tasks. Compared with the baseline damage-cost setting, the average relative cost advantage remains similar on training tasks but increases on generalization tasks.

**Figure S4:**
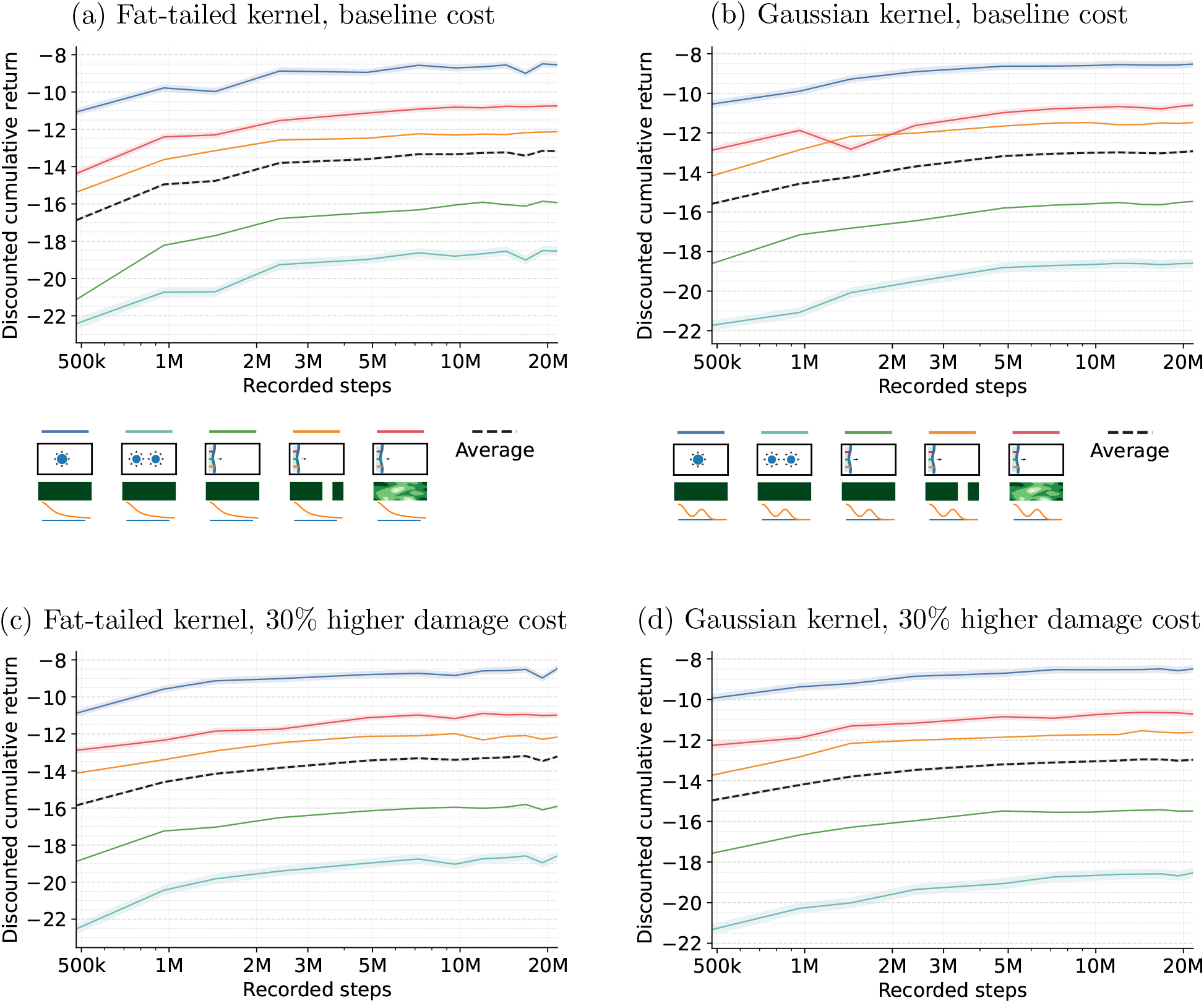
Learning progress across dispersal kernels and damage-cost settings. Solid curves show the mean discounted cumulative return for each training task at checkpoints through 21.6 million training steps on the 10 × 40 grid. The dashed curve shows the equal-weight average of these means across the five training tasks. Shaded bands indicate one standard error. Return is the negative of discounted cumulative cost, so higher returns indicate lower costs. Colors identify invasion-pattern and host-landscape combinations, as shown in the shared legend (see also Figure 6 **C**). Columns correspond to fat-tailed (left) and Gaussian (right) dispersal kernels; rows correspond to baseline (top) and 30% higher (bottom) damage costs.

## References

[1] Christin, S., Hervet, É. & Lecomte, N. Applications for deep learning in ecology. Methods in Ecology and Evolution 10, 1632–1644 (2019).

[2] Pichler, M. & Hartig, F. Machine learning and deep learning—a review for ecologists. Methods in Ecology and Evolution 14, 994–1016 (2023).

[3] Simberloff, D. et al. Impacts of biological invasions: What’s what and the way forward. Trends in Ecology & Evolution 28, 58–66 (2013).

[4] Pyšek, P. et al. Scientists’ warning on invasive alien species. Biological Reviews 95, 1511–1534 (2020).

[5] IPBES. Thematic assessment report on invasive alien species and their control of the intergovernmental science-policy platform on biodiversity and ecosystem services. Tech. Rep., IPBES secretariat, Bonn, Germany (2023).

[6] Diagne, C. et al. High and rising economic costs of biological invasions worldwide. Nature 592, 571–576 (2021).

[7] Moody, M. E. & Mack, R. N. Controlling the spread of plant invasions: The importance of nascent foci. Journal of Applied Ecology 25, 1009–1021 (1988).

[8] Shigesada, N., Kawasaki, K. & Takeda, Y. Modeling stratified diffusion in biological invasions. The American Naturalist 146, 229–251 (1995).

[9] Hastings, A. et al. The spatial spread of invasions: new developments in theory and evidence. Ecology Letters 8, 91–101 (2005).

[10] Leung, B. et al. An ounce of prevention or a pound of cure: bioeconomic risk analysis of invasive species. Proceedings of the Royal Society of London. Series B: Biological Sciences 269, 2407–2413 (2002).

[11] Mehta, S. V., Haight, R. G., Homans, F. R., Polasky, S. & Venette, R. C. Optimal detection and control strategies for invasive species management. Ecological Economics 61, 237–245 (2007).

[12] Bogich, T. L., Liebhold, A. M. & Shea, K. To sample or eradicate? a cost minimization model for monitoring and managing an invasive species. Journal of Applied Ecology 45, 1134–1142 (2008).

[13] Hauser, C. E. & McCarthy, M. A. Streamlining ‘search and destroy’: cost-effective surveillance for invasive species management. Ecology Letters 12, 683–692 (2009).

[14] Sharov, A. A., Liebhold, A. M. & Roberts, E. A. Optimizing the use of barrier zones to slow the spread of gypsy moth (lepidoptera: Lymantriidae) in north america. Journal of Economic Entomology 91, 165–174 (1998).

[15] Sharov, A. A. & Liebhold, A. M. Bioeconomics of managing the spread of exotic pest species with barrier zones. Ecological Applications 8, 833–845 (1998).

[16] Epanchin-Niell, R. S. & Hastings, A. Controlling established invaders: Integrating economics and spread dynamics to determine optimal management. Ecology Letters 13, 528–541 (2010).

[17] Epanchin-Niell, R. S. & Wilen, J. E. Optimal spatial control of biological invasions. Journal of Environmental Economics and Management 63, 260–270 (2012).

[18] Chadès, I. et al. General rules for managing and surveying networks of pests, diseases, and endangered species. Proceedings of the National Academy of Sciences of the United States of America 108, 8323–8328 (2011).

[19] Baker, C. M. & Bode, M. Placing invasive species management in a spatiotemporal context. Ecological Applications 26, 712–725 (2016).

[20] Epanchin-Niell, R. S. Economics of invasive species policy and management. Biological Invasions 19, 3333–3354 (2017).

[21] Büyüktahtakın, İ. E. & Haight, R. G. A review of operations research models in invasive species management: state of the art, challenges, and future directions. Annals of Operations Research 271, 357–403 (2018).

[22] Hall, K. M., Albers, H. J., Alkaee Taleghan, M. & Dietterich, T. G. Optimal spatial-dynamic management of stochastic species invasions. Environmental and Resource Economics 70, 403–427 (2018).

[23] Nicol, S., Sabbadin, R., Peyrard, N. & Chadès, I. Finding the best management policy to eradicate invasive species from spatial ecological networks with simultaneous actions. Journal of Applied Ecology 54, 1989–1999 (2017).

[24] Chalak, M., Polyakov, M. & Pannell, D. J. Economics of controlling invasive species: A stochastic optimization model for a spatial-dynamic process. American Journal of Agricultural Economics 99, 123–139 (2017).

[25] Pepin, K. M. et al. Optimizing management of invasions in an uncertain world using dynamic spatial models. Ecological Applications 32, e2628 (2022).

[26] Bertsekas, D. P. & Tsitsiklis, J. N. Neuro-Dynamic Programming (Athena Scientific, Belmont, MA, 1996).

[27] Sutton, R. S. & Barto, A. G. Reinforcement Learning: An Introduction (MIT Press, Cambridge, MA, 1998).

[28] Fonnesbeck, C. J. Solving dynamic wildlife resource optimization problems using reinforcement learning. Natural Resource Modeling 18, 1–40 (2005).

[29] Dietterich, T., Alkaee Taleghan, M. & Crowley, M. PAC optimal planning for invasive species management: Improved exploration for reinforcement learning from simulator-defined MDPs. Proceedings of the AAAI Conference on Artificial Intelligence 27, 1270–1276 (2013).

[30] Khosravi, M. et al. AgGym: An agricultural biotic stress simulation environment for ultra-precision management planning. Computers and Electronics in Agriculture 246, 111479 (2026). URL 10.1016/j.compag.2026.111479.

[31] Chapman, M., Xu, L., Lapeyrolerie, M. & Boettiger, C. Bridging adaptive management and reinforcement learning for more robust decisions. Philosophical Transactions of the Royal Society B 378, 20220195 (2023).

[32] Mnih, V. et al. Human-level control through deep reinforcement learning. Nature 518, 529–533 (2015).

[33] Lapeyrolerie, M., Chapman, M. S., Norman, K. E. A. & Boettiger, C. Deep reinforcement learning for conservation decisions. Methods in Ecology and Evolution 13, 2649–2662 (2022).

[34] Silvestro, D., Goria, S., Sterner, T. & Antonelli, A. Improving biodiversity protection through artificial intelligence. Nature Sustainability 5, 415–424 (2022).

[35] Equihua, J., Beckmann, M. & Seppelt, R. Connectivity conservation planning through deep reinforcement learning. Methods in Ecology and Evolution 15, 779–790 (2024).

[36] Tavakoli, A., Pardo, F. & Kormushev, P. Action branching architectures for deep reinforcement learning. Proceedings of the AAAI Conference on Artificial Intelligence 32, 4131–4138 (2018).

[37] Delarue, A., Anderson, R. & Tjandraatmadja, C. Reinforcement learning with combinatorial actions: An application to vehicle routing. In Advances in Neural Information Processing Systems, vol. 33, 609–620 (2020). URL https://proceedings.neurips.cc/paper/2020/hash/06a9d51e04213572ef0720dd27a84792-Abstract.html.

[38] Dulac-Arnold, G. et al. Deep reinforcement learning in large discrete action spaces. arXiv preprint arXiv:1512.07679 (2015). 1512.07679.

[39] Lovett, G. M. et al. Nonnative forest insects and pathogens in the United States: Impacts and policy options. Ecological Applications 26, 1437–1455 (2016).

[40] Tobin, P. C. & Blackburn, L. M. (eds.) Slow the Spread: a national program to manage the gypsy moth. No. NRS-6 in General Technical Report (U.S. Department of Agriculture, Forest Service, Northern Research Station, Newtown Square, PA, 2007).

[41] Coleman, T. W. & Liebhold, A. M. (eds.) Slow the spread: a 20-year reflection on the national Lymantria dispar integrated pest management program. No. NRS-212 in General Technical Report (U.S. Department of Agriculture, Forest Service, Northern Research Station, Madison, WI, 2023).

[42] Vercken, E., Kramer, A. M., Tobin, P. C. & Drake, J. M. Critical patch size generated by Allee effect in gypsy moth, Lymantria dispar (l.). Ecology Letters 14, 179–186 (2011).

[43] Lampert, A. & Liebhold, A. M. Optimizing the use of suppression zones for containment of invasive species. Ecological Applications 33, e2797 (2023).

[44] Lampert, A. Optimizing strategies for slowing the spread of invasive species. PLOS Computational Biology 20, e1011996 (2024).

[45] Lampert, A. & Liebhold, A. M. Optimal strategies for utilizing host plant distributions to slow the spread of plant pests. Journal of Applied Ecology 63, e70312 (2026).

[46] Liebhold, A. M. & Kean, J. M. Eradication and containment of non-native forest insects: successes and failures. Journal of Pest Science 92, 83–91 (2019).

[47] Tobin, P. C. & Blackburn, L. M. Long-distance dispersal of the gypsy moth (Lepidoptera: Lymantriidae) facilitated its initial invasion of Wisconsin. Environmental Entomology 37, 87–93 (2008).

[48] Coleman, T. W., Foelker, C. J. & Perkins, T. A 20-year synthesis of the Lymantria dispar Slow the Spread program. In Coleman, T. W. & Liebhold, A. M. (eds.) Slow the spread: a 20-year reflection on the national Lymantria dispar integrated pest management program, no. NRS-212 in General Technical Report, chap. 3, 45–69 (U.S. Department of Agriculture, Forest Service, Northern Research Station, Madison, WI, 2023).

[49] Tobin, P. C. Cost analysis and biological ramifications for implementing the gypsy moth slow the spread program. General Technical Report NRS-37, U.S. Department of Agriculture, Forest Service, Northern Research Station, Newtown Square, PA (2008).

[50] Dulac-Arnold, G. et al. Challenges of real-world reinforcement learning: definitions, benchmarks and analysis. Machine Learning 110, 2419–2468 (2021).

[51] Barto, A. G., Sutton, R. S. & Anderson, C. W. Neuronlike adaptive elements that can solve difficult learning control problems. IEEE Transactions on Systems, Man, and Cybernetics SMC-13, 834–846 (1983).

[52] Anthony, T., Tian, Z. & Barber, D. Thinking fast and slow with deep learning and tree search. In Advances in Neural Information Processing Systems, vol. 30, 5360–5370 (2017).

[53] Silver, D. et al. Mastering the game of Go without human knowledge. Nature 550, 354–359 (2017).

[54] Stern, V. M., Smith, R. F., van den Bosch, R. & Hagen, K. S. The integration of chemical and biological control of the spotted alfalfa aphid: The integrated control concept. Hilgardia 29, 81–101 (1959).

[55] Kogan, M. Integrated pest management: Historical perspectives and contemporary developments. Annual Review of Entomology 43, 243–270 (1998).

[56] Johnson, D. M., Liebhold, A. M., Tobin, P. C. & Bjørnstad, O. N. Allee effects and pulsed invasion by the gypsy moth. Nature 444, 361–363 (2006).

[57] Walter, J. A., Firebaugh, A. L., Tobin, P. C. & Haynes, K. J. Invasion in patchy landscapes is affected by dispersal mortality and mate-finding failure. Ecology 97, 3389–3401 (2016).

[58] Silvestro, D. et al. Using artificial intelligence to optimize ecological restoration for climate and biodiversity. bioRxiv (2025). URL https://www.biorxiv.org/content/10.1101/2025.01.31.635975v2. Preprint.

[59] Lauer, C. J., Montgomery, C. A. & Dietterich, T. G. Spatial interactions and optimal forest management on a fire-threatened landscape. Forest Policy and Economics 83, 107–120 (2017).

[60] García, J. & Fernández, F. A comprehensive survey on safe reinforcement learning. Journal of Machine Learning Research 16, 1437–1480 (2015). URL https://www.jmlr.org/papers/v16/garcia15a.html.

[61] McDonald-Madden, E. et al. Monitoring does not always count. Trends in Ecology & Evolution 25, 547–550 (2010).

[62] Lampert, A. & Liebhold, A. M. Combining multiple tactics over time for cost-effective eradication of invading insect populations. Ecology Letters 24, 279–287 (2021).

[63] Buczkowski, G. & Wossler, T. C. Controlling invasive Argentine ants, Linepithema humile, in conservation areas using horizontal insecticide transfer. Scientific Reports 9, 19495 (2019). URL 10.1038/s41598-019-56189-1.

[64] Thorne, B. L., Alspach, S., Tenn, K. E. & Clark, M. S. Actions toward containment, control, and eradication of invasive conehead termites (Blattodea: Isoptera: Termitidae). Journal of Integrated Pest Management 12, 22 (2021). URL 10.1093/jipm/pmab015.

[65] Pluess, T. et al. Which factors affect the success or failure of eradication campaigns against alien species? PLOS ONE 7, e48157 (2012). URL 10.1371/journal.pone.0048157.

[66] Tobin, P. C. et al. Determinants of successful arthropod eradication programs. Biological Invasions 16, 401–414 (2014). URL 10.1007/s10530-013-0529-5.

[67] Blackwood, J. C. et al. Bioeconomic synergy between tactics for insect eradication in the presence of Allee effects. Proceedings of the Royal Society B: Biological Sciences 279, 2807–2815 (2012). URL 10.1098/rspb.2012.0255.

[68] Schaul, T., Quan, J., Antonoglou, I. & Silver, D. Prioritized experience replay. In International Conference on Learning Representations (ICLR) (2016). URL https://arxiv.org/abs/1511.05952. ArXiv:1511.05952.

[69] Lillicrap, T. P. et al. Continuous control with deep reinforcement learning. In 4th International Conference on Learning Representations (ICLR 2016), Conference Track Proceedings (San Juan, Puerto Rico, 2016). URL https://arxiv.org/abs/1509.02971.1509.02971.

[70] He, K., Zhang, X., Ren, S. & Sun, J. Deep residual learning for image recognition. In 2016 IEEE Conference on Computer Vision and Pattern Recognition (CVPR), 770–778 (2016).

[71] Ha, Y., Shashaani, S. & Tran-Dinh, Q. Improved complexity of trust-region optimization for zeroth-order stochastic oracles with adaptive sampling. In 2021 Winter Simulation Conference (WSC), 1–12 (2021).

[72] Ha, Y. & Shashaani, S. Iteration complexity and finite-time efficiency of adaptive sampling trust-region methods for stochastic derivative-free optimization. IISE Transactions 57, 541–555 (2025).

[73] Eckman, D., Henderson, S., Shashaani, S. & Pasupathy, R. SimOpt (2025). URL https://simopt.org. Version 1.2.2.

[74] Kandasamy, K. et al. Tuning hyperparameters without grad students: Scalable and robust bayesian optimisation with dragonfly. Journal of Machine Learning Research 21, 1–27 (2020). URL http://jmlr.org/papers/v21/18-223.html.

